# Rhodanese Rdl2 produces reactive sulfur species to scavenge hydroxyl radical and protect mitochondria

**DOI:** 10.1101/2021.07.19.452856

**Authors:** Qingda Wang, Zhigang Chen, Xi Zhang, Yuping Xin, Yongzhen Xia, Luying Xun, Huaiwei Liu

**Affiliations:** State Key Laboratory of Microbial Technology, Shandong University, Qingdao, 266237, PR China; Department of Chemistry, School of Molecular Biosciences, Washington State University, Pullman, WA, 99164-4630, USA

**Author notes:** These authors contribute equally to this work (Qingda Wang, Zhigang Chen). Correspondence: Smith Center 519, Washington State University, Pullman, WA, 99164-7520, USA.; 72 Binhai Road, Qingdao, 266237, People’s Republic of China.

**Keywords:** Rhodanese, antioxidant, reactive sulfane sulfur, Fenton reaction, hydroxyl radical, mitochondria health

## Abstract

During aerobic respiration, mitochondria generate superoxide anion (O_2_^•−^), hydrogen peroxide (H_2_O_2_), and hydroxyl radical (HO^•^), and these reactive oxygen species (ROS) are detrimental to mitochondria. Mitochondrial damage is linked to a broad spectrum of pathologies such as Alzheimer’s disease, hemochromatosis, and diabetes. Mitochondria contain several enzymes for rapidly removing superoxide anion and hydrogen peroxide, but how they antagonize HO^•^ is elusive, representing a loophole in the anti-ROS system. Herein, we discovered that Rhodanese 2 (Rdl2) is critical for maintaining the functionality and integrity of mitochondria under sub-lethal ROS stress in *Saccharomyces cerevisiae*. Rdl2 converts stable sulfur species (thiosulfate and dialkyl polysulfide) to reactive sulfane sulfur including persulfide that protects mitochondrial DNA via scavenging HO^•^. Surprisingly, hydrogen sulfide (H_2_S) promotes HO^•^ production through stimulating the Fenton reaction, leading to increased DNA damage. Our study may reveal an *ex-ante* mean for antagonizing HO^•^, patching the loophole of the anti-ROS system in mitochondria.

## Introduction

Mitochondria are essential organelles in eukaryotic cells. They produce about 80% of cellular energy via oxidative phosphorylation under oxic condition [1]. Further, they have other key functions including heme production, iron-sulfur cluster biogenesis, and calcium homeostasis [2]. Maintaining the functionality and integrity of mitochondria is critical for human health, and failing to do so would lead to a broad spectrum of pathologies, such as Alzheimer’s, Parkinson’s disease, hemochromatosis, atherosclerosis, carcinogenesis, and diabetes mellitus [3–6].

As a hectic distribution station of metabolites, mitochondria incessantly experience many types of stresses. A widely documented one is mediated by reactive oxygen species (ROS) that includes the superoxide anion (O_2_^•−^), the hydrogen peroxide (H_2_O_2_) and the hydroxyl radical (HO^•^) [7]. O_2_^•−^ is directly produced by Complex I, III, and IV via transferring one electron to O_2_. The scavenging of O_2_^•−^ is conducted by superoxide dismutases (SOD), which convert O_2_^•−^ to H_2_O_2_. GPX enzymes including peroxiredoxins, thioredoxins, and glutathione peroxidases are the major H_2_O_2_ scavengers [7,8]. SOD and GPX compose a major anti ROS system in mitochondria. Albeit the existence of these enzymes, a fraction of H_2_O_2_ still converts to HO^•^ via the Fenton reaction. Different from H_2_O_2_, HO^•^ is a very strong and labile oxidant (2.33 V, at pH 7) that rapidly reacts with most organic molecules, such as DNA, proteins, lipids, and polysaccharides, causing structural damages to mitochondria if out of control. It is commonly believed that mitochondria only have *ex-post* means for antagonizing HO^•^—resorting to the repair systems for mending the damages [9], and no a specific enzyme or compound has been assigned for specifically eliminating it. Therefore, how mitochondria maintain HO^•^ under lethal level is a loophole in their anti ROS system.

In recently years, a new type of cellular compounds defined as reactive sulfane sulfur (RSS) has been discovered in both eukaryotic and prokaryotic cells [10–13]. RSS includes inorganic polysulfide (HS_n_H, n≥2), organic polysulfide (RS_n_H, n≥2), and polysulfane (RS_n_R, n≥2) [14]. Although several RSS generation enzymes including cystathionine gamma-lyase, cystathionine beta-synthase, and 3-mercaptopyruvate sulfurtransferase (Mst) have been identified in cytoplasm [15,16], mitochondria are thought to be the main RSS generation organelles in mammals, where sulfide:quinone oxidoreductase (Sqr) oxidizes hydrogen sulfide (H_2_S) to RSS and excessive RSS is oxidized by persulfide dioxygenase (Pdo) to sulfite [17]. Cysteinyl-tRNA synthetase 2 (Crs2) also has unspecific RSS generation function via converting cysteine to cysteine persulfide (CysSSH) in mitochondria [18]. Impairing RSS biogenesis leads to mitochondrial dysfunction, intimating a close connection between RSS biogenesis and mitochondrial health, but the underlying mechanism remains elusive [19,20].

*Saccharomyces cerevisiae* mitochondria have no Sqr or Pdo. In 2019, Nishimura et al. discovered that *S. cerevisiae* cysteinyl-tRNA synthetase (Crs1) is expressed via an alternative transcriptional initiation to locate into mitochondria, and hence it may pertain to mitochondrial RSS generation [21]. Recently, we found that rhodanese 1 and 2 (Rdl1 and Rdl2), members of the sulfurtransferase family, are responsible for thiosulfate assimilation in *S. cerevisiae*, and during this process, glutathione persulfide (GSSH) is formed as an intermediate [22]. Herein, we verified that rhodanese 2 (Rdl2) is the main RSS generation enzyme in mitochondria and more importantly, it is crucial for maintaining mitochondrial health under ROS stress. Knocking out Rdl2 results in the change of mitochondrial morphology, loss of mitochondrial DNA (mitDNA), reduction of oxidative phosphorylation efficiency, and change of iron metabolism. RSS generated by Rdl2 actively scavenges HO^•^. Thus, an active mechanism to remove HO^•^ might be identified.

## Materials and Methods

### Strains and materials

*S. cerevisiae* BY4742 (*MATα his3Δ1 leu2Δ0 lys2Δ0 ura3Δ0*) and CEN.PK2 (MATa/MATα *ura3-52/ura3-52; trp1-289/trp1-289; leu2-3,112/leu2-3,112; his3Δ 1/his3Δ 1; MAL2-8C/MAL2-8C; SUC2/SUC2*) strains were cultured in yeast extract-peptone-dextrose (YPD) medium or synthetic defined (SD) medium at 30°C. The constructed mutants are listed in supplementary material (Table S1). *E. coli* DH5α and BL21(DE3) strains were cultured in lysogeny broth (LB) medium at 37°C. Thiosulfate and sodium hydrosulfide (NaHS) were purchased from Sigma-Aldrich (Saint Louis, MO). Dimethyl trisulfide (MeSSSMe) and S_8_ were purchased from TCI (Shanghai, China) Company. HS_n_H and GSSH were prepared following the protocol of Luebke et al [23]. Other chemicals were purchased from local companies if not specifically mentioned.

Key methods were described as below and others were provided in supplementary material (Materials and methods).

### Subcellular Localization of Rdl1 and Rdl2

A GFP encoding sequence was fused to *RDL1* and *RDL2* C-termini in BY4742 chromosome by using the one-step PCR-mediated gene disruption method [24]. Yeast cells were collected by centrifugation (10,000g, 5 min) and diluted to 1 OD_600_ in sterile SD medium containing 10 nM Mito Tracker Red CMSRos (Thermo Fisher, Waltham, MA). The cell suspensions were incubated at 30°C for 20 min in dark place and then washed three times to remove the extracellular CMSRos. Yeast cells were suspended in water and imaged by using a fluorescence microscope (Olympus IX83). GFP and CMSRos co-localization was analyzed.

To analyze the expression level of fused Rdl2-GFP in different growth phages, Microplate Reader Synergy H1 was used with *λ_ex_* of 488 nm and *λ_em_* of 525 nm. To analyze the expression level of fused Rdl2-GFP in the presence of H_2_O_2_, 1 OD_600_ middle-log phase yeast cells (8 h cultivation) were treated with 5 mM H_2_O_2_ at 30°C for 1 h, and then subjected to flow cytometry analysis. For each sample, 10^5^ cells were analyzed in the FL1 channel and the average fluorescence value was recorded.

### Transcriptomic analysis

*S. cerevisiae* strains were cultured in YPG medium until OD_600_ reached 0.8, and then 2□mM□ □H_2_O_2_ was added and the cultivation was expanded for 12 h. Cells were harvested for the omics analysis, which were performed at Shanghai Applied Protein Technology Co., Ltd (Shanghai, China). For transcriptomic analysis, total RNA was extracted. Magnetic beads with Oligo (dT) were used to enrich mRNA and fragmentation buffer was added to randomly interrupt the mRNA. The first strand of cDNA was obtained with six-base random primers and the second strand of cDNA was synthesized by adding buffer, dNTPs and DNA polymerase I. Double-stranded cDNA was purified with AMPure XP and then A-tailing and sequencing adapters were connected. The AMPure XP beads were used for fragment size selection and PCR enrichment was perform to obtain the final cDNA library. The library was sequenced on Illumina NovaSeq 6000 platform. Sequencing was performed at Shanghai Applied Protein Technology Co., Ltd. The clean data were obtained from raw data by removing reads containing adapter, poly-N and low quality reads. The clean reads were aligned with the genome of *S. cerevisiae* BY4742 by using HISAT2. The featureCounts software was used to calculate the FPKM value of each gene expression in each sample. Genes with a p-vale<0.05 and fold change>2 were considered as significantly differentially expressed.

### Targeted metabolomics analysis

For targeted metabolomics analysis, cells were harvested and quickly frozen in liquid nitrogen. After grinding with liquid nitrogen, 60 mg cells were mixed with 1 ml methanol acetonitrile aqueous solution (2:2:1, v/v) and vortexed for 60 s. Cells were then disrupted using the ultra-sound method. The broken cells were placed at −20 °C for 1 h and then centrifuged at 14,000g for 20 min at 4°C. The obtained supernatant was subjected to freeze-dry and then LC-MS analysis. The Waters I-class ultra performance liquid chromatography was used and the mobile phase A was an aqueous solution with 25 mM ammonium acetate and 25 mM ammonia (pH, 9.75). Mobile phase B was acetonitrile. Protein sample was placed in a 4°C autosampler, and the column temperature was set to 40°C. The flow rate was 0.3 mL/min, and the injection volume was 2 μL. The liquid phase gradient was set as: 0-1 min, phase B at 95%; 1-14 min, B linearly changing from 95% to 65%; 14-16 min, B linearly changing from 65% to 40%; 16-18 min, B at 40%; 18-18.1 min, B linearly changing from 40% to 95%; 18.1-23 min, B at 95%. In the sample cohort, one QC sample was set every six experimental repeats to detect and evaluate the stability and repeatability of the system.

The AB 5500 QqQ mass spectrometer (AB SCIEX, Framingham, MA) was sued for mass spectrometry analysis. The ESI source conditions were as: sheath gas temperature, 350°C; dry gas temperature, 350°C; sheath gas flow, 11 L/min; dry gas flow, 10 L/min; capillary voltage was 4000 V for positive mode and −3500 V for negative mode; nozzle voltage, 500 V; and nebulizer pressure, 30 psi. Monitor was in MRM mode and the dwell time of each MRM transition was 3 ms, and the total cycle time wa 1.263 s. MRMAnalyzer (R) was used to extract the original MRM data of 200 metabolites to obtain the peak area of each metabolite. Metabolites with p-value<0.05 and fold change>1.5 were considered as at significantly different levels.

### Mitochondria preparation and mitochondrial RSS analysis

*S. cerevisiae* mitochondria were isolated using differential centrifugation as described previously [25]. The fluorescence-based probe psGFP1.1 was expressed and localized into mitochondrial matrix of BY4742 as reported previously [26]. Mitochondria containing psGFP1.1 (mit-psGFP) were isolated and treated with sulfur containing chemicals in isolation buffer (1 mM EDTA, 0.6 M sorbitol, 10 mM Tris-HCl, pH 7.4). The mixtures contained mitochondria and 400 μM cysteine, thiosulfate, or MeSSSMe. After incubated at 30°C for 1 h, treated mitochondria were centrifuged at 12,000g for 15 min and washed with isolation buffer (1 mM EDTA, 0.6 M sorbitol, 10 mM Tris-HCl, pH 7.4). Fluorescence was detected using the Synergy H1 microplate reader. The emission intensities at 515 nm excited by both 408 nm and 488 nm were recorded, and the ratio of 408/488 was used to represent the reactive sulfane sulfur level in mitochondria. Explanation of the calculation principle can be found in a previous report [26].

### Radical-induced pDNA cleavage assay

The pDNA cleavage assay was performed following the previously reported method [27]. The pBluescript II SK (+) plasmid was used as the model DNA. 25 μg/μL plasmid was mixed with 50 μM Fe^2+^ and 50 μM H_2_O_2_ in distilled water. The mixture was incubated at room temperature for 3 h and then analyzed by electrophoresis. The supercoiled (SC) and nicked circular (NC) forms of plasmid was quantified using the software FluorChem Q (Protein Simple, Inc.). SC ratio was calculated as SC/(SC+NC) and used as the DNA damage index. For testing the protection effect of Rdl2 products, Rdl2 (2.5 μM) was incubated with 1 mM thiosulfate or MeSSSMe at room temperature for 1 h, and then Rdl2 was removed using a 3-kDa filter. The filtrated products were diluted with HEPES buffer (100 mM, pH 7.4) to prepare product-dilution solutions, which were added into the DNA-Fe^2+^-H_2_O_2_ mixture and subjected to the same analysis. Thiosulfate, MeSSSMe, H_2_S (in the form of NaHS), S_8_, laboratory prepared GSSH, GSH, and L-ascorbic acid were also tested in the same conditions.

For testing whether H_2_S reduces Fe^3+^ to promote the Fenton reaction, 25 μg/μL plasmid was incubated with 50 μM Fe^3+^, 50 μM H_2_O_2_ and NaHS or S_8_ (20 μM-300 μM) at room temperature for 3 h. As the control, 25 μg/μL plasmid was incubated with 50 μM H_2_O_2_ and NaHS (20 μM-300 μM) at room temperature for 3 h. The untreated-plasmid DNA and the Fenton solution-plasmid DNA mixture were also used as controls.

### In vitro analysis of produced H_2_S and reactive sulfane sulfur

A modified methylene blue method [28] was used to detect the produced H_2_S. Briefly, 1% zinc acetate solution (900 μL) was added into the reaction solution (100 μL) to convert H_2_S to ZnS. The mixture was centrifuged at 8,000g for 5 min, and washed with deionized and distilled water. The washing step was repeated three times. After removing the final supernatants, deionized and distilled water (100 μL) was added to the precipitates (ZnS) and adequately mixed. Then, the solution was incubated with 300 μL of 1% zinc acetate, 50 μL of 20 mM DPDA (N,N-dimethyl-p-phenylenediamine) and 30 mM FeCl_3_ in 7.2 N HCl for 30 min. The samples were centrifuged at 8,000g for 5 min and transferred into 96-well plates to measure the OD_630_. NaHS was used to make the standard curve.

Reactive sulfane sulfur was analyzed by using a previously reported method [29]. Briefly, the sample (50 μL) was mixed with 25 mM monobromobimane (mBBr, 5 μL) in acetonitrile and incubated in the dark at room temperature for 30 min. An equal volume of acetic acid and acetonitrile mixture (v/v, 1:9) was added to precipitate proteins. The precipitates were removed via centrifugation at 12,000 *g* for 2 min. The obtained supernatant was subjected to LC-ESI-MS analysis (Ultimate 3000, Burker impact HD).

### Statistical Analysis

Transcriptomics and targeted metabolomics analysis were performed with six parallel biological samples. The data have been deposited in https://www.biosino.org/node/ with ID: OEP001468. Other analysis were performed with at least 3 parallel biological samples. Data are presented as mean ± S.D.

## Results

### Rdl2 is the main enzyme for RSS biogenesis in S. cerevisiae mitochondria

To check if Crs1 was mainly responsible for generating RSS in yeast mitochondria, we expressed it in *Escherichia coli* BL21(DE3) and purified it. The enzymatic activity was examined by mixing Crs1 with PLP (pyridoxal phosphate, the cofactor) and cysteine (the substrate). The product was analyzed by using the sulfane sulfur prober 4 (SSP4) [30] and LC-MS, and only trace amounts of cysteine persulfide (Cys-SSH) were produced (Fig. 1A and supplementary material, Figure S1). Crs1 was overexpressed in *S. cerevisiae* BY4742 from two different promoters, the inducible promoter *P_gal_* and the constitutive promoter *P_tef1_*; neither one led to increased RSS production *in vivo* (Fig. 1B and 1C). Since knocking out Crs1 in haploid strain BY4742 is lethal, we performed the knock-out experiment in the diploid strain CEN.PK2. The Crs1 knock-out strain (*crs1*^-/+^) showed no detectable difference from its wild type (wt) in terms of intracellular RSS level (Fig. 1D). The above experiments suggested that Crs1’s contribution to the mitochondrial RSS is limited.

**Fig. 1.**
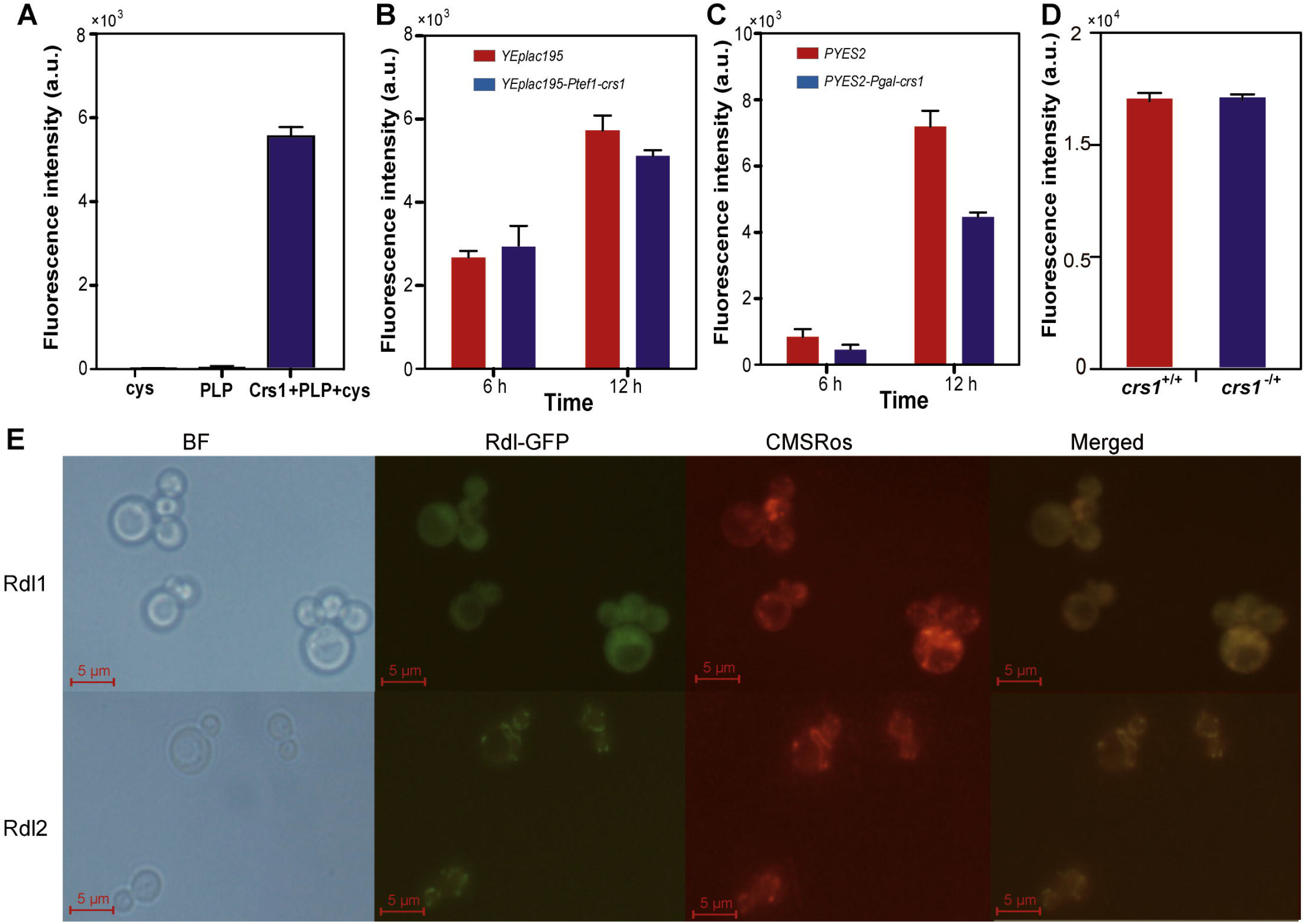
Activity and localization analyses of the enzymes pertaining to mitochondrial RSS generation in *S. cerevisiae* BY4742. A) SSP4 analysis of RSS generated by Crs1 *in vitro*. B&C) SSP4 analysis of intracellular RSS of BY4742 containing the *opo* plasmid (as control) and the Crs1 expression plasmid. D) SSP4 analysis of intracellular RSS of the CEN CEN.PK2 wt (*crs1^+/+^*) and *CRS1* knock-out (*crs1*^-/+^) strains. E) Localization analysis of Rdl1 and 2. CMSRos is the Mito Tracker Red dye. Data were from three independent repeats and represented as average ± s.d.

To check whether Rdl1 and Rdl2 were mainly responsible for RSS generation in yeast mitochondria, we first investigated the locations of these two enzymes. A green fluorescent protein (GFP) was fused with Rdl1 and Rdl2 by inserting the GFP ORF into the C-termini of Rdl1 and 2 ORFs in *S. cerevisiae* BY4742 genome. The Mito Tracker Red CMSRos was used to label BY4742 mitochondria. Rdl1/2-GFP and CMSRos co-location analysis showed that Rdl2 particularly located in mitochondria; whereas, Rdl1 mainly located in cytoplasm (Fig. 1E). We isolated intact mitochondria from BY4742 (wt) and the *RDL2* deletion (*Δrdl2*) mutant, in which the mitochondrial RSS detection probe mit-psGFP [26] was expressed. The wt mitochondria contained more RSS than did the *Δrdl2* counterparts, as represented by a higher 408/488 fluorescence ratio (Fig. 2A). The isolated mitochondria were then treated with cysteine, thiosulfate, or organic polysulfide (RS_n_R, n≥2)—the three potential substrates of sulfurtransferases. The addition of cysteine caused no RSS increase, indicating that again Crs1 did not use cysteine to produce RSS. Thiosulfate and dimethyl trisulfide (MeSSSMe) caused 27% and 261% increases of RSS, respectively, in wt mitochondria (Fig. 2B). Whereas, thiosulfate caused no increase, but MeSSSMe caused 66% increase of RSS in *Δrdl2* mitochondria (Fig. 2C). The *RDL2* gene was cloned and expressed in *E. coli* BL21 (DE3). The addition of thiosulfate and MeSSSMe increased RSS in *RDL2*-expressing *E. coli*, but did not in the *E. coli* host (Fig. 2D and 2E).

**Fig. 2.**
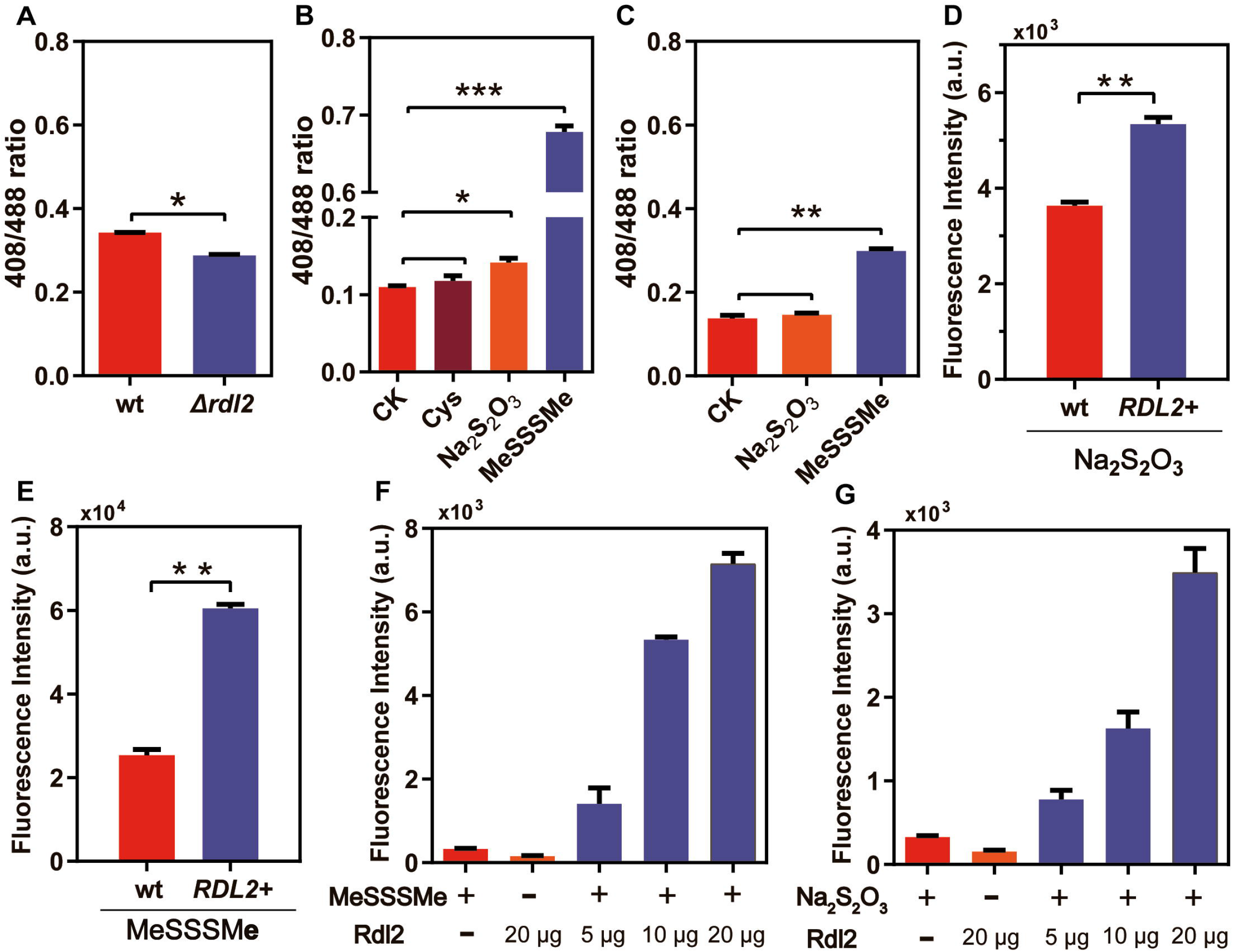
*In vitro* and *in vivo* analysis of the Rdl2 contribution to mitochondrial RSS. A) BY4742 wt and *Δrdl2* strains harboring mit-psGFP were cultured in YPD medium until OD600 reached 0.6, and then 408/488 excitation ratio of the cells was examined. B) Mitochondria harboring mit-psGFP were isolated from BY4742 wt strain, and were treated with 400 μM different chemicals at 30°C for 1 h, and then 408/488 excitation ratio was examined. The higher the ratio, the higher the RSS. C) Mitochondria harboring mit-psGFP were isolated from BY4742 *Δrdl2* strain, and were treated with 400 μM different chemicals at 30°C for 1 h, and then 408/488 excitation ratio was examined. D&E) *E. coli* wt and *rdl2* expressing strains were treated with thiosulfate and MeSSSMe. SSP4 was used to to examine the intracellular reactive sulfane sulfur. F&G) Purified Rdl2 was mixed with its substrates and the products were analyzed using SSP4. Data were from three independent repeats. * represents difference (p<0.05) and ** represents significant difference (p<0.01) in two-sided t-test. Data were from three independent repeats and represented as average ± s.d

RSS production by Rdl2 was further tested with purified Rdl2. It produced RSS from thiosulfate and MeSSSMe as detected with SSP4; the amount of RSS was positively correlated to the amount of added Rdl2 (Fig. 2F and 2G). No apparent production of H_2_S was detected by using a modified methylene blue method [28]. When the products of MeSSSMe reacting with Rdl2 were derivatized by monobromobimane (mBBr) and analyzed by using LC-MS. A major peak corresponding to mB-SS-mB was detected (supplementary material, Figure S2A), indicating the production of hydrogen persulfide (HS_2_H, HS_2_^-^, or S_2_^2-^). A much smaller peak of likely mB-SSS-mB was also observed (supplementary material, Figure S2B). MeSSMe, but neither MeSMe nor Me-SS-mB, was detected (supplementary material, Figure S2C), indicating that Rdl2 releases a sulfane sulfur atom from MeSSSMe to produce HS_n_H and MeSSMe.

### Rdl2 is essential for maintaining mitochondrial health under sub-lethal levels of ROS

Compared with wt strain, the *Δrdl2* strain showed slightly growth retardation on both a fermentable carbon source (glucose) and a none-fermentable carbon source (glycerol). When a sub-lethal level of H_2_O_2_ was present, the *Δrdl2* strain displayed more severe growth retardation than wt strain, especially on none-fermentable carbon source (Fig. 3A-3C), suggesting that mitochondria were impaired in the *Δrdl2* strain.

**Fig. 3.**
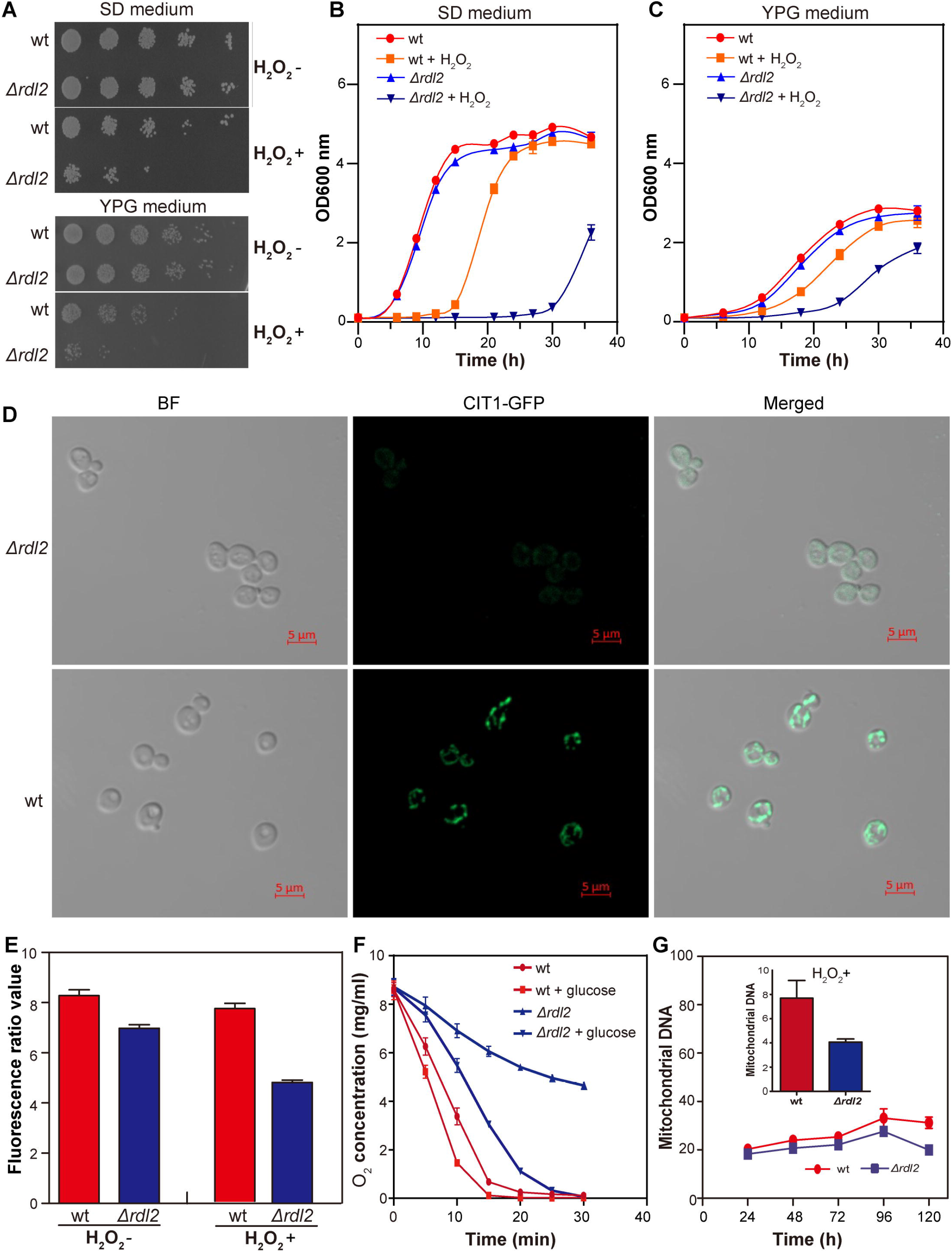
Examination of the role of Rdl2 in maintaining mitochondrial health under ROS stress. A) BY4742 wt and *Δrdl2* strains were cultivated in SD or YPG agar plates. 2 mM H_2_O_2_ was added. B&C) BY4742 wt and *Δrdl2* strains were cultivated in SD or YPG liquid medium. 2 mM H_2_O_2_ was added. D) Mitochondrial morphology analysis of BY4742 wt and *Δrdl2* strains. The strains containing Cit1-GFP were cultivated in YPD medium to log phase without H_2_O_2_ treatment. Images were captured with the laser confocal microscope LMS900. E) Mitochondrial membrane potential analysis of wt and *Δrdl2* strains. F) Oxygen consumption analysis of wt and *Δrdl2* strains. G) mit-DNA analysis of wt and *Δrdl2* strains. Embedded figure: the strains were treated with 2 mM H_2_O_2_. Data were calculated from three independent repeats.

A GFP ORF was fused with the mitochondrial citrate synthase 1 (Cit1) encoding gene in genomes of both wt and *Δrdl2* strains, which were cultured in YPD medium without the addition of H_2_O_2_. The mitochondrial morphology was observed with a laser-scanning confocal microscope. Mitochondria in wt displayed normal morphology with mainly filamentous or granular shape; whereas, mitochondria in the *Δrdl2* strain displayed irregular shape (Fig. 3D). The GFP fluorescence intensity was also obviously lower in *Δrdl2* than in wt. These phenomena indicated that mitochondria in the *Δrdl2* strain were abnormal.

Several mitochondria-related physiological characteristics were then examined. First, the mitochondrial membrane potential of *Δrdl2* was lower than that of wt no matter with or without the addition of H_2_O_2_ (Fig. 3E). Second, the oxygen consumption rate of the *Δrdl2* strain was obviously lower than that of wt with or without the presence of glucose (Fig. 3F). Third, the relative number of mitDNA (mitochondrial DNA normalized against nuclear DNA) was lower in *Δrdl2* than in wt, especially in the presence of H_2_O_2_ (Fig. 3G). These results verified that the health of mitochondria was impaired by Rdl2 knock-out, and the impairment made *Δrdl2* more sensitive to ROS stress.

### Systematically investigation of the physiological changes caused by knocking out Rdl2

Transcriptomics and metabolomics approaches were applied to systemically analyze the consequences of Rdl2 knock-out. Both wt and *Δrdl2* strains were treated with 2 mM H_2_O_2_ because their physiological differences were more obvious under ROS stress. In transcriptome level, 215 genes were downregulated and 156 genes were upregulated (fold change >2, p<0.05) in the *Δrdl2* strain compared to wt (Fig. 4A). For specification:

I. Among the 117 genes involved in glycolysis/gluconeogenesis/pentose phosphate pathway, only 11 of them were changed at transcription level (Fig. 4B), indicating the metabolism in cytoplasm was not severely affected by Rdl2 deletion. However, or 10 genes involved in tricarboxylic acid (TCA) cycle, 7 were downregulated and 3 were upregulated (Fig. 4C); for 20 genes involved in fatty acid degradation, 8 of them were downregulated (supplementary material, Figure S3); for 11 genes involved in respiration chain, 9 of them were downregulated and 2 were upregulated (Fig. 4D);
II. For 13 genes involved in anti-ROS processes, 9 of them were downregulated and 4 were upregulated (Fig. 4E); for 38 genes involved in responding to DNA damage and replication stress, 23 of them were downregulated and 15 were upregulated; for 4 genes involved in maintaining iron homeostasis, 1 of them were downregulated and 3 were upregulated (Fig. 4F).

**Fig. 4.**
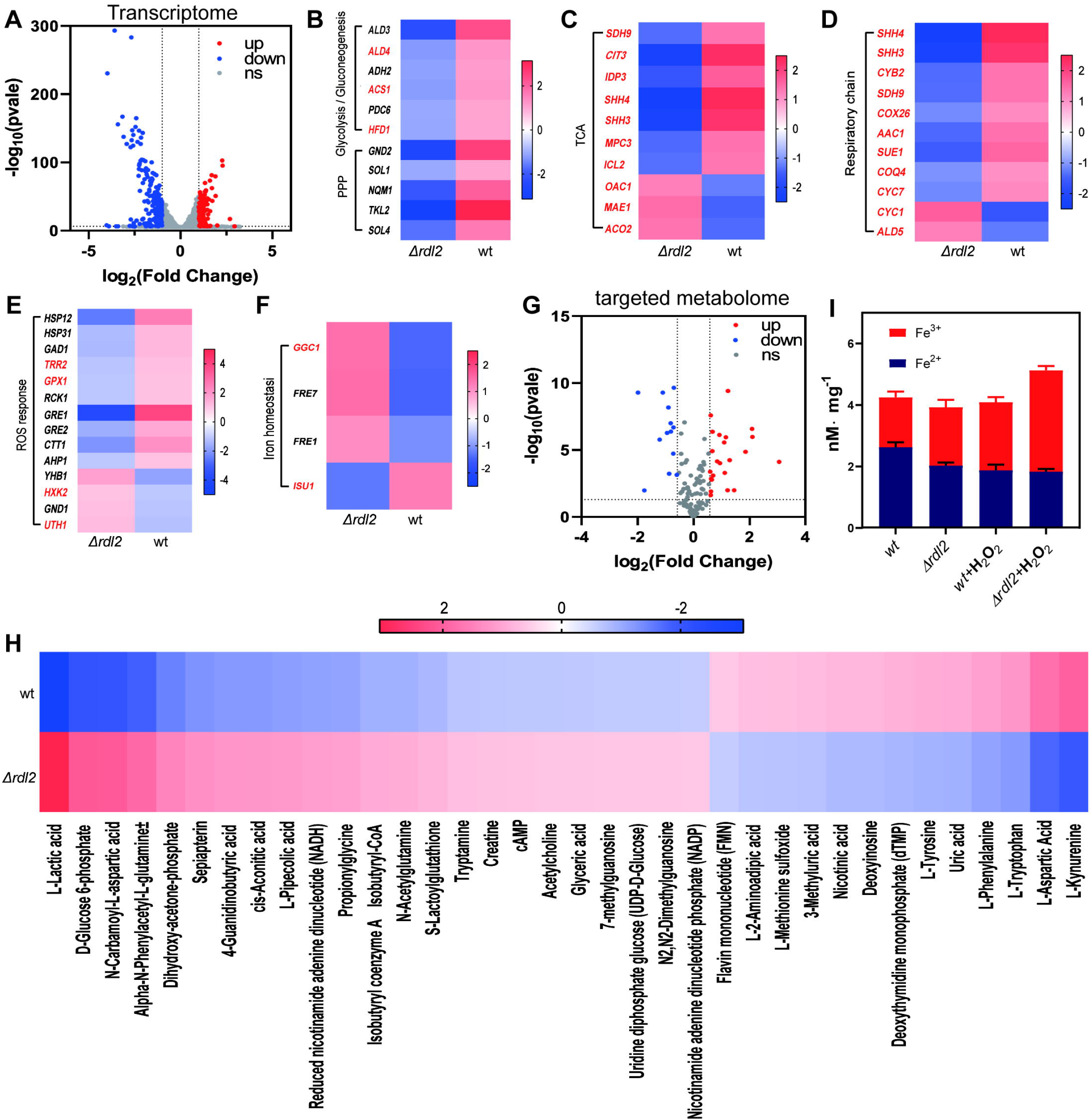
Transcriptomics and targeted metabolomics analysis of BY4742 wt and *Δrdl2* strains under the stress of 2 mM H_2_O_2_. A-F) Genes changed at transcriptional level. G&H) Concentration changed metabolites. I) intracellular iron analysis. Transcriptomics and targeted metabolomics analysis were performed with six parallel biological samples. Iron analysis data were calculated from three independent repeats.

Together, these results indicated that the energy generation processes that mainly happen in mitochondria were impaired, and some anti-DNA damage processes were activated by Rdl2 knock-out. It is noteworthy that *OAC1*, the gene involved in TCA cycle but also responsible for thiosulfate importing into mitochondria was downregulated, implying a less thiosulfate import to mitochondria. The well-known anti-ROS genes including mitochondrial thioredoxin reductase (*TRR1*), thiol-specific peroxiredoxin (*AHP1*), and cytosolic catalase (*CTT1*) were all downregulated in *Δrdl2* strain, implying that their anti-ROS functions are not over-lapped with Rdl2 (otherwise, they should be upregulated to compensate for the loss of Rdl2 function). The upregulation of plasma membrane ferric importing/reduction enzymes (*FRE7* and *FRE1*) and the mitochondrial iron transporter (*GGC1*), together with the downregulation of mitochondrial iron-sulfur assembly enzyme (*ISU1*, which consumes reduced iron) implied that there was a lack of reduced iron (Fe^2+^) in mitochondria of *Δrdl2* strain.

For metabolomics analysis, we targeted 200 metabolites of central and energetic metabolism (supplementary material, sheet 1). Among them, 131 compounds were identified with 36 showing significant abundance change between wt and *Δrdl2* strains (fold change >1.5, p<0.05) (Fig. 4G). For specification:

I. The abundance of 23 metabolites was increased in *Δrdl2* strain compared to that of wt strain (Fig. 4H). The increase of reduced nicotinamide adenine dinucleotide (NADH) and lactic acid indicated that the consumption of NADH through the respiration chain was reduced.
II. The abundance of 13 metabolites was decreased in the *Δrdl2* strain compared to that of wt strain, including flavin mononucleotide (FMN), the main cofactor of enzymes composing the respiration chain (Fig. 4H).

The metabolome included no iron, we performed iron analysis additionally. Without ROS stress, the *Δrdl2* strain contained less Fe^2+^ and more Fe^3+^ compared to wt strain. The addition of H_2_O_2_ increased the content of Fe^3+^ and decreased the content of Fe^2+^ in wt strain (Fig. 4I). In *Δrdl2* strain, the content of Fe^3+^ was increased but content of Fe^2+^ was not by H_2_O_2_, indicating that more Fe^3+^ was imported into the cell (Fig. 4I). These phenomena were in consistent with the results of transcriptomic analysis.

### Rdl2-generated RSS protects DNA from hydroxyl radial-induced damage

We suspected that the Rdl2-generated RSS may protect mitDNA via interfering with the Fenton reaction. To test this hypothesis, the classical radical-induced plasmid DNA (pDNA) cleavage assay [27] was performed. The principle of this assay is when Fe^2+^, H_2_O_2_, and pDNA are mixed in deionized water, HO^•^ generated from the Fenton reaction cleaves the supercoiled plasmids (SC) and converts most of them to nicked-circular form (NC), but when a HO^•^ or H_2_O_2_ scavenger is present, the damage to pDNA is inhibited. Fe^2+^ (50 μM), H_2_O_2_ (50 μM), and pDNA (25 μg/μl) were used. We first tested thiosulfate and MeSSSMe (20 μM ~ 300 μM), and observed no protection of SC by either one. Second, we tested the products of Rdl2 + thiosulfate and Rdl2 + MeSSSMe reactions, SC percentage was apparently increased by each one, indicating a pDNA protection effect (Fig. 5A). As two controls, laboratory prepared glutathione persulfide (GSSH, made via mixing GSH with S_8_) and S_8_ also showed SC protection effect (Fig. 5B and 5C). These results indicated that RSS indeed interfere with the Fenton reaction.

**Fig. 5.**
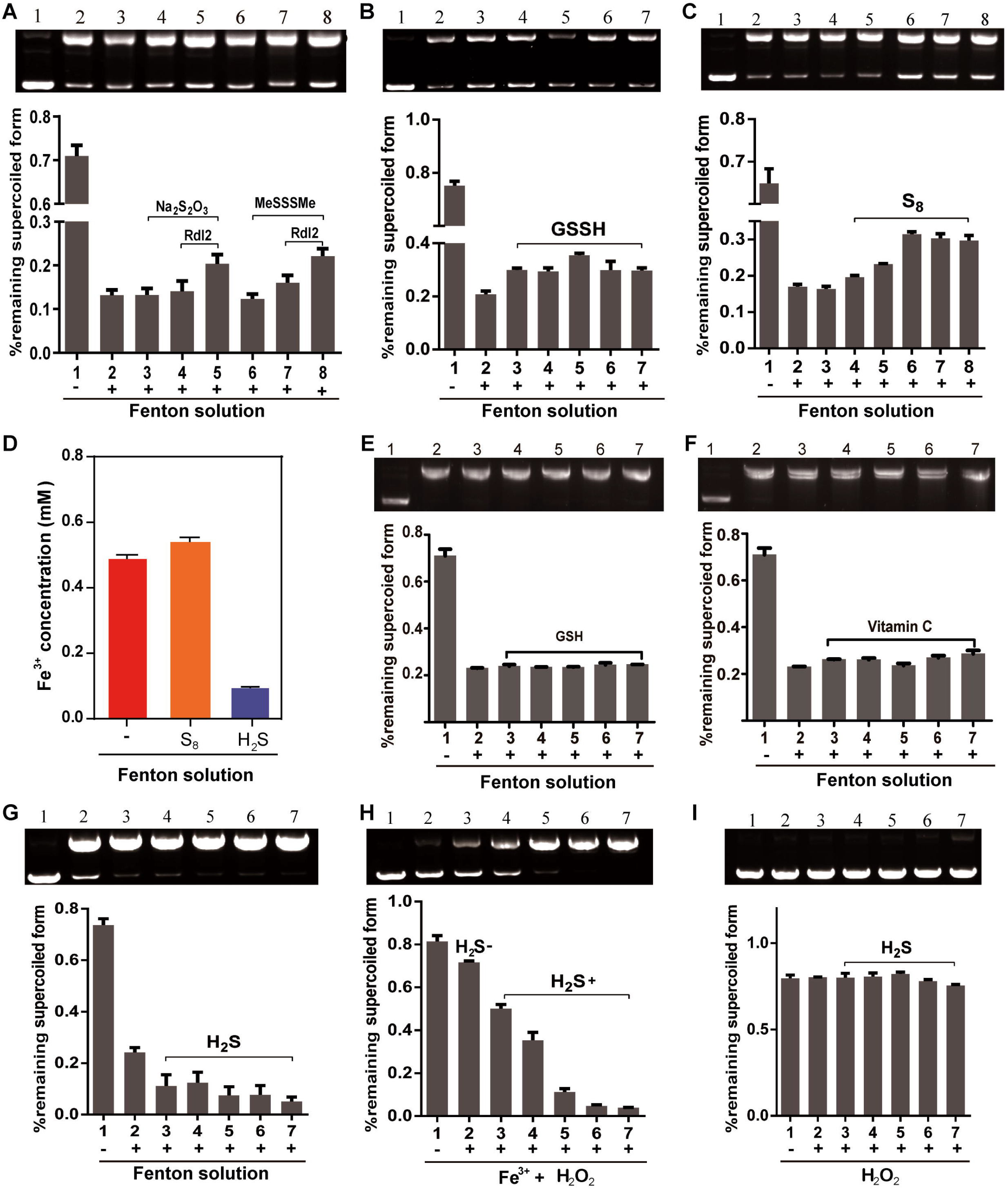
Plasmid DNA cleavage and Fenton reaction assays performed with Rdl2 products and different chemicals. Upper figure shows the gel electrophoresis results of pDNA and lower figure shows the percentage of pDNA in SC form. A) 1, untreated pDNA; 2, pDNA treated with Fenton reagents (50 μM Fe^2+^ and 50 μM H_2_O_2_); 3, pDNA treated with Fenton reagents and thiosulfate (300 μM); 4, pDNA treated with Fenton reagents and products from Rdl2 + thiosulfate reaction (10 fold-dilution); 5, pDNA treated with Fenton reagents and products from Rdl2 + thiosulfate reaction (2.5 fold dilution); 6, pDNA treated with Fenton reagents and MeSSSMe (300 μM); 7, pDNA treated with Fenton reagents and products from Rdl2 + MeSSSMe reaction (10 fold-dilution); 8, pDNA treated with Fenton reagents and products from Rdl2 + MeSSSMe reaction (2.5 fold-dilution). B,C,E,F,&G) 1, untreated pDNA; 2, pDNA treated with Fenton reagents; 3-n, pDNA treated with Fenton reagents and tested chemicals (20 μM-300 μM). D) Fe^3+^ production from the Fenton reaction. CK is the mixture of 500 μM Fe^2+^ + 500 μM H_2_O_2_; 500 μM S_8_ or H_2_S was added. H) 1, untreated pDNA; 2, pDNA treated with 50 μM Fe^3+^ + 50 μM H_2_O_2_; 3-7, pDNA treated with 50 μM Fe^3+^, 50 μM H_2_O_2_, and H_2_S (20 μM-300 μM). I) 1, untreated pDNA; 2, pDNA treated with 50 μM H_2_O_2_; 3-7, pDNA treated with 50 μM H_2_O_2_ and H_2_S (20 μM-300 μM). The SC percentage data were calculated from three independent repeats.

RSS may function through reacting with H_2_O_2_ to decrease the production of HO^•^, or scavenging the produced HO^•^. We examined the Fe^3+^ production from the Fenton reaction with or without the addition of S_8_. The Fe^3+^ production from the Fe^2+^ + H_2_O_2_ + S_8_ solution was not decreased but slightly higher than that without S_8_ (Fig. 5D), suggesting that S_8_ did not react with H_2_O_2_ in the Fenton solution. The slight increase in Fe^3+^ production in the presence of S_8_ favorably argues that S_8_ reacts HO^•^ and drives the Fenton reaction forward. We also determined the rate constant of the S_8_ + H_2_O_2_ reaction to be 1.04 M^−1^·s^−1^ (supplementary material, Figure S4), much lower than the reported rate constant of the Fenton reaction, ~10^3^ M^−1^·s^−1^ [31], further supporting that S_8_ does not react with H_2_O_2_ in the Fenton solution. GSH and L-ascobic acid are well-known cellular antioxidants [32,33], but neither of them showed SC DNA protection at similar concentrations to RSS (20 μM ~ 300 μM, Fig. 5E and 5F).Thus, S_8_ likely scavenges HO^•^ to protect SC.

H_2_S was recognized as an endogenous antioxidant in a profound study that led the tide of H_2_S related research [34]. However, we found that it promoted pDNA cleavage in the Fenton solution (Fig. 5G), indicating it stimulates the Fenton reaction. To reveal the underlying mechanism, we mixed Fe^3+^ (instead of Fe^2+^), H_2_O_2_, and pDNA with or without H_2_S. The pDNA cleavage was observed in the presence of H_2_S, but not in its absence (Fig. 5H). Without iron, the H_2_O_2_ solution with H_2_S showed no pDNA cleavage (Fig. 5I). These results suggested that H_2_S functions through reducing Fe^3+^ to Fe^2+^ to facilitate the Fenton reaction. For verification, we analyzed the Fe^3+^ production from the Fenton reaction with or without H_2_S, and much higher Fe^3+^ was detected from the Fenton reaction without H_2_S than with H_2_S (Fig. 5D). Black precipitate (FeS) was temporary produced in the Fe^2+^ + H_2_O_2_ + H_2_S solution, but it disappeared in 2 min, indicating that S^2−^ in FeS is still available for reacting with H_2_O_2_ and Fe^3+^. These results suggest that H_2_S enhances the Fenton reaction.

We then measured the rate constants of sulfur-containing compounds reacting with HO^•^. H_2_S slightly descended the absorbance ratio (A_0_/A) of the HO^•^ probe, suggesting that it actually enhanced the production of HO^•^ (from Fenton reaction) other than scavenging it (Fig. 6A). GSH modestly reacted with HO^•^, represented by a relative flat curve of the absorbance ratio; whereas GSSH and S_8_ rapidly reacted with HO^•^ with 13.32×10^9^ and 8.15×10^9^ rate constants, respectively (Fig. 6B). These results, again, indicated that RSS was efficient HO^•^ scavenger.

**Fig. 6.**
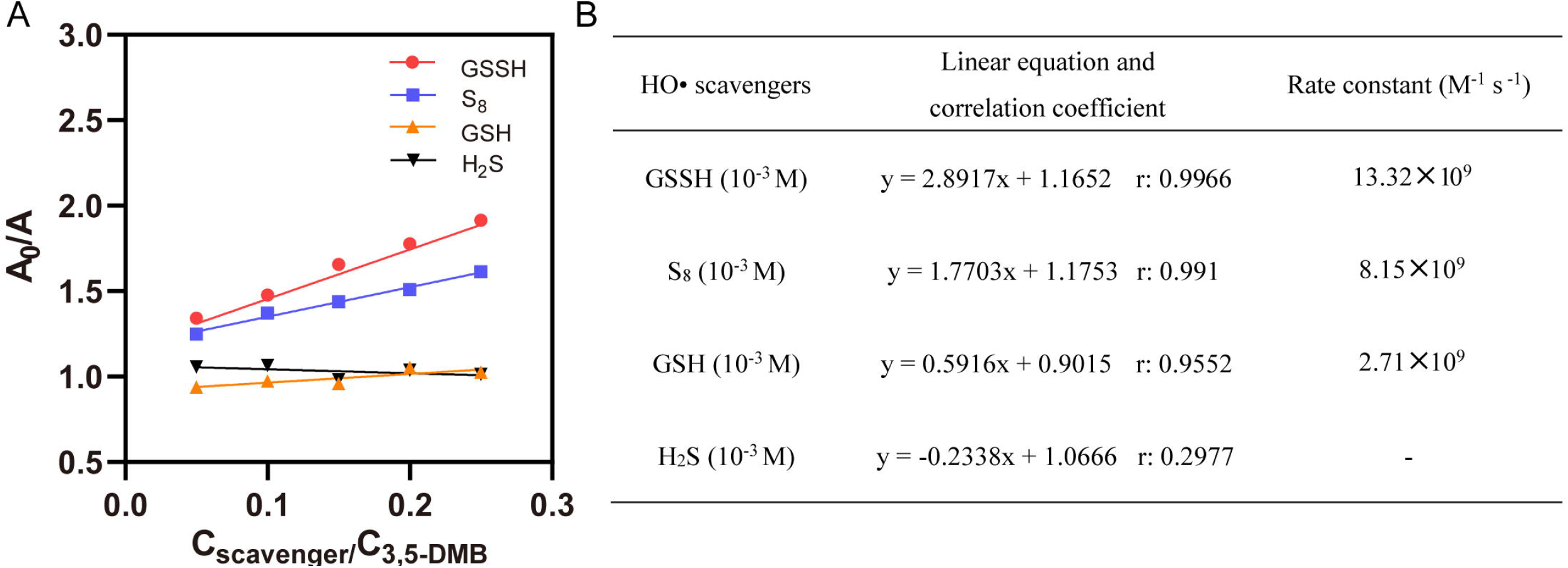
Kinetic analysis of HO^•^ radical scavenging activity of certain compounds using the modified CUPRAC method. A) Kinetic plots measured using the HO^•^ probe 3,5-dimethoxybenzoate. B) Rate constants calculated from the kinetic plots. Details of the CUPRAC method was described in supporting material.

### RDL2 expresses and functions in post-logarithmic phase

The *Δrdl2* strain released less H_2_S than wt strain, detected by lead acetate papers (Fig. 7A). We checked the transcription levels (obtained from the transcriptome data) of enzymes involved in H_2_S production pathways. Cbs (encoding gene is *CYS3*) and Cse (encoding gene is *CYS4*) that are responsible for H_2_S production from organic substrate (mainly cysteine) were up-regulated in *Δrdl2* strain (Fig. 7B). The enzymes responsible for H_2_S production from inorganic substrates (sulfate and sulfite) were not significantly changed. These results suggested that *Δrdl2* strain actually has a higher H_2_S productivity than wt strain, but most of the produced H_2_S may be consumed in Fe^3+^ reduction, leading to a less release portion.

**Fig. 7.**
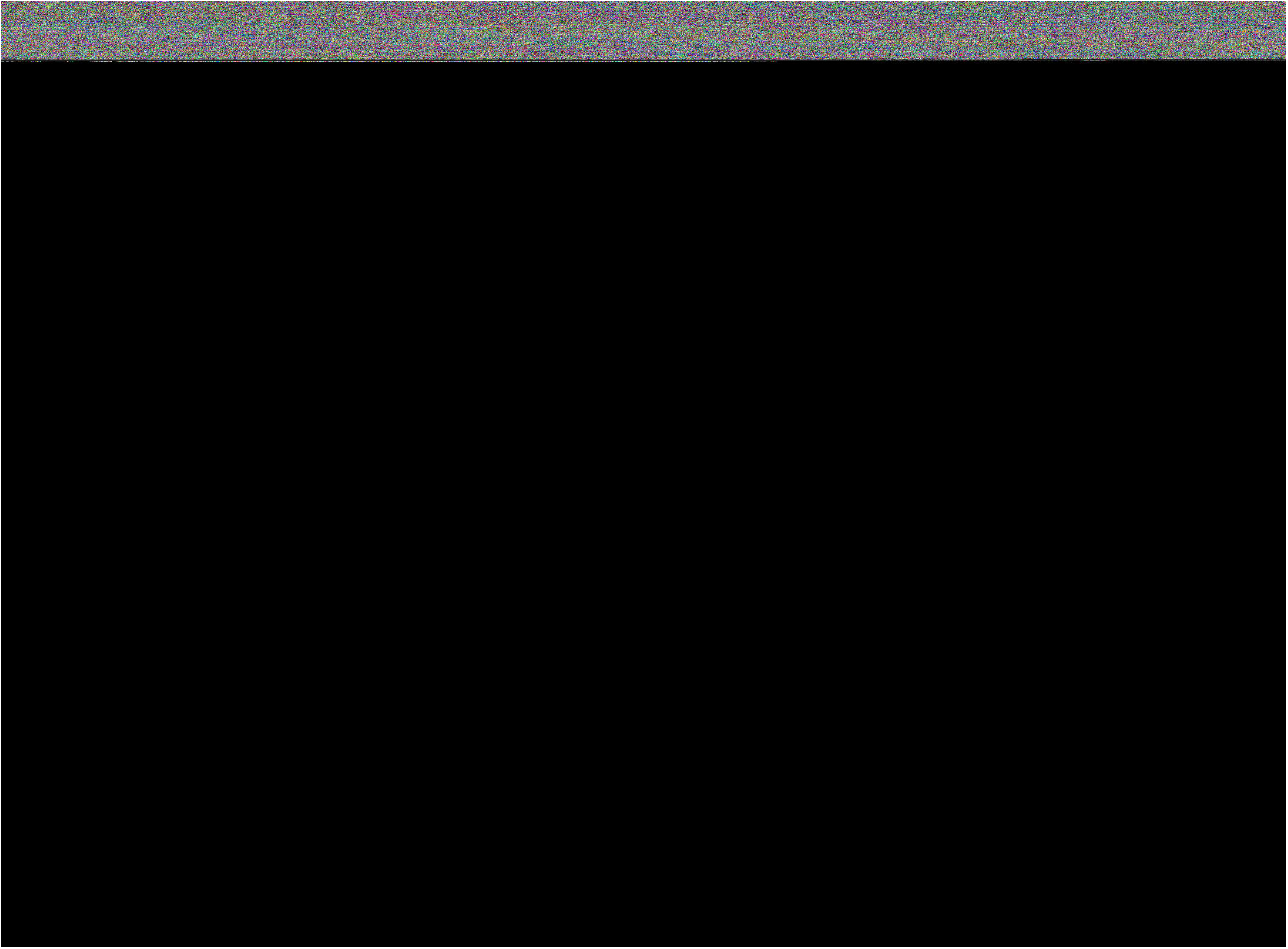
*In vitro* analysis of the interaction between Rdl2 and ROS. A) H_2_S release detection using lead acetate papers. The strains were cultivated in SD medium for 80 h. B) Transcription levels of enzymes involved in H_2_S production pathways. Data were from transcriptomic analysis. The bar represent the average readcount of 6 parallel biological samples. C) Analysis of the intracellular ROS of BY4742 wt and *Δrdl2* strains. D) The expression of Rdl2-GFP was only slightly increased in high H_2_O_2_ condition (5 mM). E) The expression of Rdl2-GFP was different at different growth phases. F) BY4742 wt cells of stationary phase were more resistant to H_2_O_2_ than mid-log phase cells. Data were from three independent repeats and represented as average ± s.d

The intracellular ROS levels of wt and *Δrdl2* strains were compared using the DCFH-DA probe. Wt strain contained constantly lower ROS level than *Δrdl2* strain and both showed increasing ROS levels with growth time expanding (Fig. 7C). When H_2_O_2_ was added, ROS level in wt strain was moderately increased; whereas, ROS level in *Δrdl2* strain was sharply increased (Fig. 7C), indicating that RSS is critical for controlling the level of intracellular ROS.

The Rdl2-GFP expressing strain was treated with H_2_O_2_, and the expression of Rdl2-GFP was only slightly increased in high H_2_O_2_ condition (> 5 mM/OD_600_ cell) (Fig. 7D), indicating that the expression of Rdl2 might not be ROS-induced. We analyzed the expression level of Rdl2-GFP in different growth periods by flow cytometry, and observed that Rdl2 mainly expressed in post-logarithmic phases (after 12 h in YPD medium) (Fig. 7E). For confirmation, we inoculated cells of 8 h-culture (in which Rdl2 had not been expressed) onto the H_2_O_2_-containing agar medium, wt cells barely grew and *Δrdl2* cells totally did not grow. When cells of 24 h-culture were inoculated (in which Rdl2 had been expressed), wt cells grew much better while *Δrdl2* cells also showed little growth (Fig. 7F). These results indicated that the expression of Rdl2 is mainly controlled by growth related system(s) and its expression is concurrent with the accumulation of intracellular ROS.

## Discussion

In this study, we verified that Rdl2 is the main RSS-generating enzyme in *S. cerevisiae* mitochondria. Rdl2 deletion resulted in RSS decrease in the mitochondria. Further, we observed that Rdl2 deletion leads to morphology change of mitochondria even without the treatment of H_2_O_2_ (Fig. 3). When facing H_2_O_2_ stress, Rdl2 deletion strain is more fragile with obvious changes in mitochondria. To understand the underlying mechanism, we studied the Rdl2 involved reactions and found that it generates RSS via releasing sulfane sulfur atoms from relatively stable sulfane sulfur carriers (Thiosulfate and RS_n_R, Fig. 2). More importantly, the RSS product of Rdl2 can scavenge HO^•^ and hence protects mitochondria from HO^•^ induced damages, such as loss of mitDNA, impair of mitochondria membrane, and disturbance of iron homeostasis.

Although we detected that HS_n_H is the main product of Rdl2 *in vitro*, considering the abundance of intracellular GSH, the main product should be GSSH *in vivo* (GSH + S_n_ → GSSH + S_n-1_). It is reported that RSSH is an excellent H-atom (H•) donor, which easily gives its H• to alkoxyl (RO^•^) with a ~10^9^ M^−1^·s^−1^ rate constant [35]. RSSH then becomes RSS^•^ radical, and two RSS^•^ radicals dimerize to form one more stable RSSSSR with a 5×10^9^ M^−1^·s^−1^ rate constant [35,36]. In contrast, the rate constant of HO^•^ reacting with DNA is at 10^8^ M^−1^·s^−1^ level, at least one magnitude lower than that of RSSH giving H-atom reaction. Thus, RS_n_H and HS_n_H chemicals have the potential capability of scavenging HO^•^ before it can damage DNA *in vivo*. For S_n_, it is unknown how it scavenges HO^•^. Possibly via donating an electron to HO^•^ and forming S_n_^•^ itself, then two S_n_^•^ radicals form S_2n_ that has a longer chain. Further studies are required to elucidate the exact mechanism. It is noteworthy that although we observed that GSH and L-ascorbic acid did not protect pDNA from the Fenton reaction even when their concentrations were 6-fold higher than that of Fenton solution (Fig. 5E and 5F), a previous study demonstrated that GSH did show DNA protection effect at much higher concentration (>25 fold of Fenton solution). The authors suggested that GSH may also react with HO^•^ through H-atom transfer mechanism [37]. Nonetheless, GSH is a much less efficient H-atom donor than RSS as verified previously [35,36].

Mitochondria antagonize ROS at three levels. The scavenging of O_2_^•−^ is conducted by SOD and the scavenging of H_2_O_2_ is performed by GPX enzymes (Fig. 7). It is reported that the rate constants of GPX reactions with H_2_O_2_ are in the range of 10^6^~10^7^ M^−1^·s^−1^ [38,39]. Metal iron mediated-Fenton reaction is the main origin of HO^•^ in mitochondria with a rate constant of ~10^3^ M^−1^·s^−1^ [31]. Considering the physiological concentrations of H_2_O_2_ and Fe^2+^ are ~20 nM and ~500 nM, respectively [8], and the rate constants of RSS + H_2_O_2_ reactions are only 1.04 M^−1^·s^−1^—23.76 M^−1^·s^−1^ [40], RSS should have no chance to react with H_2_O_2_ in mitochondria. Thus, RSS plays antioxidation function mainly through scavenging HO^•^, as we observed that it reacts with HO^•^ more rapidly than GSH. This may represent the third ROS-antagonizing level in mitochondria (Fig. 8).

**Fig. 8.**
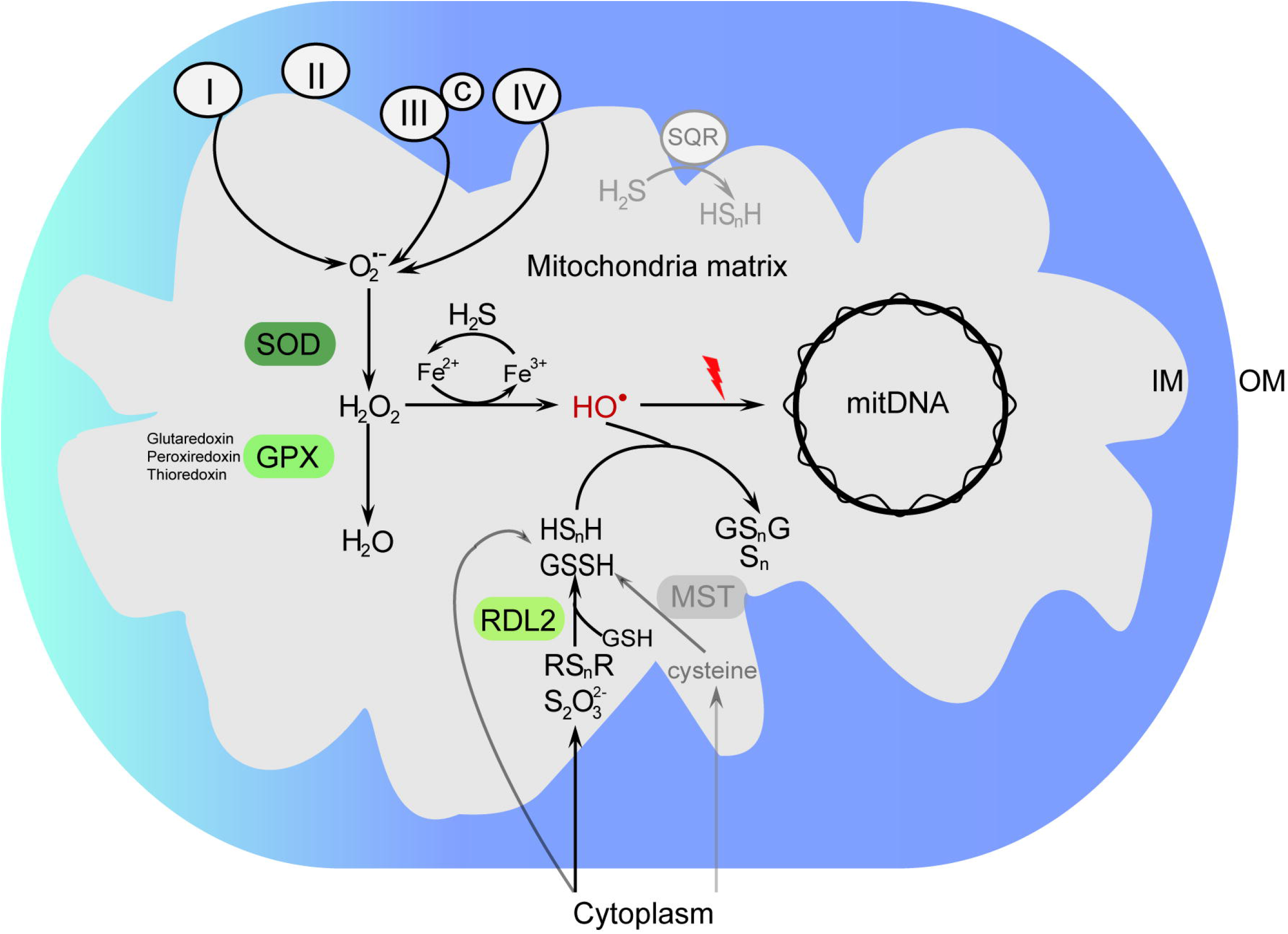
Schematic representation of the mitochondrial RSS biogenesis routes and functions. *S. cerevisiae* mitochondria contain no Sqr or Mst. Rdl2 is the only enzyme that generates RSS, which can scavenge HO^•^.

On the other hand, H_2_S has been deemed as an antioxidant in many reports [34,41,42], further studies including ours found that its reaction with H_2_O_2_ is slow with 2^nd^ rate constant between 0.46M^−1^·s^−1^—0.76 M^−1^·s^−1^ [40]. Considering the physiological concentration of H_2_S is much lower than that of RSS [15], the reaction between H_2_S and H_2_O_2_ may not happen either *in vivo*. Herein, we found that H_2_S has promoting effect on the Fenton reaction, indicating that H_2_S has the pro-oxidation function other than antioxidation.

In terms of the whole eukaryotic kingdom, there are probably four routes for mitochondria to obtain reactive sulfane sulfur: exporting from cytoplasm, generating from H_2_S via Sqr, generating from cysteine via Mst or Crs2, and generating from sulfane sulfur stocks via rhodanese (Fig. 8). These routes may function with different efficiencies and in different conditions. Sqr-mediated route might be the most efficient one; however, Sqr activity is ETC (electron transport chain) dependent, hence it may function mostly in hypoxic condition. Rhodanese-mediated route is oxygen independent, they may function in all conditions. For cells having no Sqr (such as *S. cerevisiae*) or whose Sqr is not active (such as neuron), the rhodanese-mediated RSS biogenesis represents a general strategy for protecting cells from HO^•^-induced damage.

In conclusion, our study has three important observations:

i. RSS has HO^•^ scavenging function, highlighting the unparallel role of RSS among all known natural anti ROS agents.
ii. H_2_S stimulates the Fenton reaction, suggesting that it may promote oxidative damage, instead of being an antioxidant.
iii. Rhodaneses are critical for maintaining mitochondrial health, adding a new function for them in anti ROS besides sulfur metabolism.

## Acknowledgements

The work was financially supported by grants from the National Key R&D Program of China (2018YFA0901200) and the National Natural Science Foundation of China (91951202).

## Author Contributions

H. Liu and L. Xun designed the research and made plans for the experiments; Q. Wang, Z. Chen, and Y. Xin performed Rdl2 related experiments; X. Zhang performed Crs1 related experiments. Y. Xia helped in data interpretation.

## Competing interests

The authors declare no competing interests.

## Data availability

The data that support the findings of this study are available from the corresponding author upon request

## Notes

https://www.biosino.org/node/

### Competing Interest Statement

The authors have declared no competing interest.

