## Supplementary material for "Rhodanese Rdl2 produces reactive sulfur species to scavenge hydroxyl radical and protect mitochondria"

**Contents**

Materials and methods	2

Figure S1	10

Figure S2	11

Figure S3	12

Figure S4	13

Sheet 1. List of the targeted 200 metabolites	14

References	18

### Materials and methods

*Strains and plasmids*

*Table S1. The strains and plasmids used in this study*

| Strain/plasmid | Relevant characteristic(s) | source |
| --- | --- | --- |
| ***S. cerevisiae*** |  |  |
| BY4742 | *MAT**α his3**Δ1 leu2**Δ0 lys2**Δ0 ura3Δ0* | Lab stock |
| BY4742 *Δrdl2* | *Δrdl2*::*BLE* | Lab stock |
| BY4742 *RDL2-GFP* | *RDL2-GFP(S65T)-HIS3* | This study |
| BY4742 *CIT1-GFP* | *CIT1-GFP(S65T)-HIS3* | This study |
| BY4742 *Δrdl2* *CIT1-GFP* | *Δrdl2*::*BLE CIT1-GFP(S65T)-HIS3* | This study |
| BY4742: YEplac195 | Wild type with YEplac195 | This study |
| BY4742: YEplac195-*CRS1* | Expression of *CRS1* with YEplac195 plasmid | This study |
| BY4742: mit-psGFP | Expression of psGFP in BY4742 | Lab stock |
| BY4742*Δrdl2*: mit-psGFP | Expression of psGFP in BY4742*Δrdl2* | This study |
| BY4742: pYES2 | Wild type with pYES2 plasmid | This study |
| BY4742: pYES2-*CRS1* | Expression of *CRS1* with pYES2 plasmid | This study |
| CEN.PK2 | *MATa/MATα ura3-52/ura3-52; trp1-289/trp1-289; leu2-3,112/leu2-3,112; his3Δ 1/his3Δ 1; MAL2-8C/MAL2-8C; SUC2/SUC2* | Lab stock |
| CEN.PK2 *crs1^-/+^* | *CRS1/ crs1::HIS3* | This study |
| ***E. coli*** |  |  |
| DH5*α* | *supE44 ΔlacU169(Φ80dlacZΔM15) hsdR17 recA1 endA1 gyrA96 thi-1 relA1* | Lab stock |
| BL21(DE3) | *F-ompT hsdSB (rB-mB-) gal ( λ1857 ind1 Sam7 nin5 lacUV5 T7gene1) dcm.* | Lab stock |
| **Plasmid** |  |  |
| YEplac195 | Multicopy plasmid for expression in yeast | Lab stock |
| mit-psGFP | YEplac195 containing fusing gene of mitochondrial sigal peptidea-psGFP. Under control of TEF1 promoter | Labstock^1^ |
| YEplac195-*CRS1* | *CRS1* in YEplac195, control by TEF1 promoter | This study |
| YEplac195-MitopsGFP | Mito-psGFP in YEplac195, control by TEF1 promoter | Lab stock[1] |
| pYES2-*CRS1* | *CRS1* in pYES2, control by GAL1 promoter | Lab stock |
| pFA6a-GFP(S65T)-His3MX6 | Template plasmid | Lab stock |
| pET30a | Expression plasmid in *E. coli* | Lab stock |
| pET30a-*RDL2* | *RDL2* in pET30a, control by IPTG-induced lac promoter | This study |
| pET30a-*CRS1* | *CRS1* in pET30a, control by IPTG-induced lac promoter | This study |
| pBluescript II SK (+) | Multicopy plasmid for pDNA cleavage analysis | Lab stock |

*H_2_O_2_ sensitivity assay*

The assay was performed on both fermentable (SD medium) and none-fermentable medium (yeast-peptone-glycerol, YPG) medium. Fresh BY4742 cells were inoculated into 5 ml liquid medium and cultivated overnight at 30^o^C. Then cell suspensions were diluted to OD_600_ of 0.1 with fresh medium. 5 μl of 6-fold serial dilutions were spotted on agar plates containing 2 mM H_2_O_2_. The plates were incubated at 30°C for 48 h. The assay was also performed in liquid medium. 0.1 OD_600_ cells were cultured in 20 ml of liquid medium containing 2 mM H_2_O_2_ and cultivated at 30°C with shaking. The growth was monitored with a spectrophotometer.

*Protein expression and purification*

The gene encoding Rdl2 was amplified from genomic DNA of *S. cerevisiae* BY4742. The gene encoding Crs1 was codon optimized and chemically synthesized following a previous report [2]. ORF of these genes were ligated with pET30a plasmid using the T5 exonuclease-dependent assembly method [3]. For protein expression and purification, *E. coli* BL21(DE3) strain harboring the expression plasmid (pET30a-*RDL2* or pET30a-*CRS1*) was incubated in LB medium at 25^o^C with shaking (225rpm). Kanamycin (50 μg/ml) was added. When OD_600_ reached 0.6, 0.1 mM isopropyl β-D-1-thiogalactopyranoside (IPTG) was added to induce the expression, and the temperature was decreased to 25^o^C. The cultivation was further continued for 22 h, then cells were harvested by centrifugation and re-suspended in buffer I (20 mM Tris-HCl, 0.5 M NaCl, 20 mM imidazole, pH 8.0). Cell disruption was performed using a Pressure Cell Homogeniser (SPCH-18) at 4^o^C. Cell lysate was centrifuged to remove the debris. Target proteins in supernatant were first purified by using nickelnitrilotriacetate (Ni-NTA) agarose. Obtained proteins were then passed through the size exclusion column (Superdex 200; GE Healthcare) for further purification.

*Detection of reactive sulfane sulfur in S. cerevisiae* *and* *E. coli*

For SSP4 based analysis of intracellular RSS of *E. coli*, *E. coli* BL21 (DE3) containing pET30a-*RDL2* was cultured in LB medium at 37^o^C with shaking (225rpm). When OD_600 nm_ reached 0.6, 0.1 mM IPTG was added and the cultivation was continued at 25^o^C for 4 h with shaking (225 rpm). Cells were collected and washed twice with 50 mM HEPES buffer (pH 7.4) and then incubated with 400 µM MeSSSMe or thiosulfate at 37^o^C for 1 h. After incubation, cells were washed twice and suspended in 1mL of 50 mM HEPES buffer (pH 7.4). 1 OD_600_ cells were reacted with 20 µM SSP4 and 500 µM CTAB (Cetyltrimethylammonium Bromide) at room temperature for 30 min, then cells were subjected to fluorescence analysis. The Synergy H1 microplate reader was used with the excitation wavelength setting to 482 nm and the emission wavelength setting to 515 nm.

For SSP4 based analysis of intracellular RSS of *S. cerevisiae*, BY4742 and CEN.PK2 strains were cultivated in YPD medium. For strains containing pYES2 derived plasmids, when OD_600_ reached 0.4, cells were collected, washed two times, and then transferred into yeast-peptone-galactose medium[4]. At each sampling time, 1 OD_600_ cells were collected and reacted with 20 µM SSP4 and 500 µM CTAB in 50 mM HEPES buffer (pH 7.0) at 30^o^C for 15 min, then cells were subjected to fluorescence analysis. The Synergy H1 microplate reader was used with the excitation wavelength setting to 482 nm and the emission wavelength setting to 515 nm.

*Reaction of Rdl2 with its substrates in vitro*

Rdl2 (5-20 μg) were mixed with 5 mM substrate (thiosulfate or MeSSSMe) in HEPES buffer (100 mM, pH 7.4). The reaction were conducted at room temperature for 30 min. For detection of the produced RSS, SSP4 (10 μM) was added into the mixture. The fluorescence was detected using Synergy H1 microplate reader. The excitation wavelength was set to 482 nm and the emission wavelength was set to 515 nm. As the control, mixture containing no protein was also treated and examined following the same protocol.

For analyzing the species of Rdl2 produced RSS, the reaction solution was derivatized by monobromobimane (mBBr) and subjected to LC-ESI-MS analysis (Ultimate 3000, Burker impact HD) following a reported protocol [5].

*Reaction of Crs1 with cysteine in vitro*

Crs1 (5μM) was mixed with cysteine (100 μM) and PLP (100 μM) in reaction buffer (50 μM HEPES, 25 mM KCl, 15 mM MgCl_2_, pH 7.4). The reaction were conducted at 30^o^C for 30 min. For detection of the produced RSS, SSP4 (10 μM) was added into the mixture. The fluorescence was detected using Synergy H1 microplate reader. The excitation wavelength was set to 482 nm and the emission wavelength was set to 515 nm. As the control, mixture containing no protein was also treated and examined following the same protocol.

For analyzing the species of Crs1 produced RSS, the reaction solution was derivatized by mBBr and subjected to LC-ESI-MS analysis (Ultimate 3000, Burker impact HD) following a reported protocol [5].

*Kinetic analysis of the reaction between S_8_ and H_2_O_2_*

The rate constant of S_8_ + H_2_O_2_ reaction was determined using a previously reported method [6]. Briefly, 20 μM-200 μM H_2_O_2_ was mixed with 2 μM S_8_ in deoxygenated HEPES buffer (100 mM, pH 7.4). The resonance synchronous spectroscopy (RS_2_) intensity of S_8_ (535 nm-545 nm) was scanned at 30 *s* intervals for 3 min. The *k_obs_* value was calculated by plotting the *ln_[RS2]_* value against the reaction time. The apparent 2^nd^ -order reaction rate constant *k* was calculated using the formula: *k_obs_* = *k* × *[H_2_O_2_]*.

*Detection of Fe^3+^ produced from the Fenton reaction*

Fe^3+^ was quantified using the thiocyanate method [7]. Briefly, 500 μM H_2_O_2_ was incubated with 500 μM Fe^2+^ in deionized and distilled water at room temperature for 5 min and then 10 mM potassium thiocyanate was added to react with the produced Fe^3+^. The OD_460 nm_ absorbance was measured using the Synergy H1 microplate reader. To see the influence of S_8_ or H_2_S s on the Fenton reaction, 500 μM S_8_ or H_2_S was added into the H_2_O_2_ and Fe^2+^ mixture, and Fe^3+^ was detected following the same protocol.

*Intracellular iron analysis*

BY4742 wt and *Δrdl2* strains were cultured in YPG medium until OD_600_ reached 0.8, and then 2 mM  H_2_O_2_ were added. After 12 h of treatment, cells were harvested and washed twice with sterile water. The intracellular Fe^2+^ and total iron concentrations were detected with an Iron Assay Kit-Colorimetric (Dojindo). The Fe^3+^ concentration was calculated by subtracting Fe^2+^ concentration from the total iron concentration. *S. cerevisiae* BY4742 wt and *Δrdl2* strains without H_2_O_2_ treatment were used as controls.

*Measurement of mitochondrial membrane potential of S. cerevisiae*

The mitochondrial membrane potential of yeast was measured using the fluorescent probe JC-1 (Beyotime). Cells were grown until OD_600_ reached 1.0, and 1 mL culture was treated with 2 mM H_2_O_2_ for 30 min. Cells without H_2_O_2_ treatment were used as the control. After the treatment, cells were washed twice and suspended in JC-1 work buffer, and then loaded with JC-1 probe following the instructions of manufacturer. The membrane potential was estimated by the ratio of red to green fluorescence intensity, which was measured with the Synergy H1 microplate reader.

*ROS analysis.*

Cellular ROS was measured using the DCFH-DA probe. Fresh yeast cells were inoculated into 50 ml SD medium and incubated at 30°C with shaking (220rpm). For H_2_O_2_ treatment, 2 mM H_2_O_2_ was added when OD_600_ reached 0.5. At each sampling point, cells were collected by centrifugation (10,000*g*, 5 min) and the cell suspensions were diluted to 1.0 OD_600_ in sterile SD medium with 10 μM DCFH-DA. The cell suspensions were incubated at 30^o^C for 20 min in dark, and then were washed three times to remove the extra DCFH-DA. The fluorescence was measured using the Microplate Reader Synergy H1, and λ_ex_ was set to 488 nm and λ_em_ was set to 525 nm.

*Oxygen consumption analysis*

BY4742 strains were cultured in YPD medium until OD_600_ reached 1.0. Cells were collected by centrifugation and washed two times with PBS buffer (pH 7.4). The cell density was finally adjusted to OD600 = 10 in PBS buffer (pH 7.4). The Orion RDO meter was used to detect the O_2_ in cell solution every 5 minute. For glucose treatment, 2% glucose (w/v) was added.

*Microscopic examination of mitochondria*

The citrate synthase 1 (CIT1) encoding gene was fused with a C terminal GFP encoding gene in BY4742 wt and *Δrdl2* genomes by using the one-step PCR-mediated gene disruption method [8]. The strains containing Cit1-GFP were cultivated in YPD medium to log phase, and then cells were harvested, washed twice with distilled water, and subjected to fluorescence microscopy analysis. The laser confocal microscope LMS900 was used.

*Copy number analysis of mitDNA*

The ratio of mitDNA to nuclear DNA was determined using the quantitative real-time PCR (qRT-PCR) method. The *COX1* gene of mitochondrial DNA and *ACT1* gene of nuclear DNA were chosen for this quantification. BY4742 wt and *Δrdl2* strains were cultured in YPD medium. For each sampling point, cells were collected and their genomes were extracted. qRT-PCR was conducted with the SYBR Green reaction kit. Each run included three parallels to obtain CT values of *ACT1* and *COX1*. Calculating the interpolation of CT_COX1_ and CT_ACT1_ to get the ΔCT, 2^ΔCT^ was used to represent the relative copy number of mitochondrial DNA. For the H_2_O_2_ treatment experiment, Log phase cells (OD_600_=1) were treated with 2 mM H_2_O_2_ for 1 h, and then subjected to qRT-PCR analysis.

*Activity assay of hydroxyl radical scavenging antioxidants*

The modified CUPRAC method [9] was applied to determine the rate constants of antioxidants reacting with HO^.^. In a 1 mL reaction system, we added 300 μL of phosphate buffer (pH 7.0), 100 μL of 10 mM probe (3,5-dimethoxybenzoate), 50 μL of 20 mM Na_2_-EDTA, 50 μL of 20 mM FeCl_2_ solution, (400-x) μL H_2_O, (x) μL scavenger sample solution (x varying between 50 and 250 μL) at concentration 10^-3^ M. The reaction was started by adding 100 μL of 10 mM H_2_O_2_, and the mixture was incubated for 2 h in a water bath kept at 37^o^C. The reaction was stopped by adding 50 μL of 2 M HCl. After a short vortex, the reaction product was extracted with 1ml ethylacetate (EtAc). 100 μL of 10 mM Cu (II) , 100 μL of 7.5 mM Neocuproine (Nc), 100 μL of 1 M Ammonium acetate (NH_4_Ac), 100 μL of EtAc extract, and 100 μL of EtOH were reacted for 30 minutes, the absorbance at OD_450 nm_ of the final solution was recorded using Microplate Reader Synergy H1. The second-order rate constants of the scavengers were determined with competition kinetics by means of a linear plot of A_0_/A as a function of C_scavenger_/C_probe_, where A_0_ and A are the CUPRAC absorbances of the system in the absence and presence of the scavenger, respectively, and C is the molar concentration of relevant species. The equations for rate constant calculation were described in [9]

### Figure S1

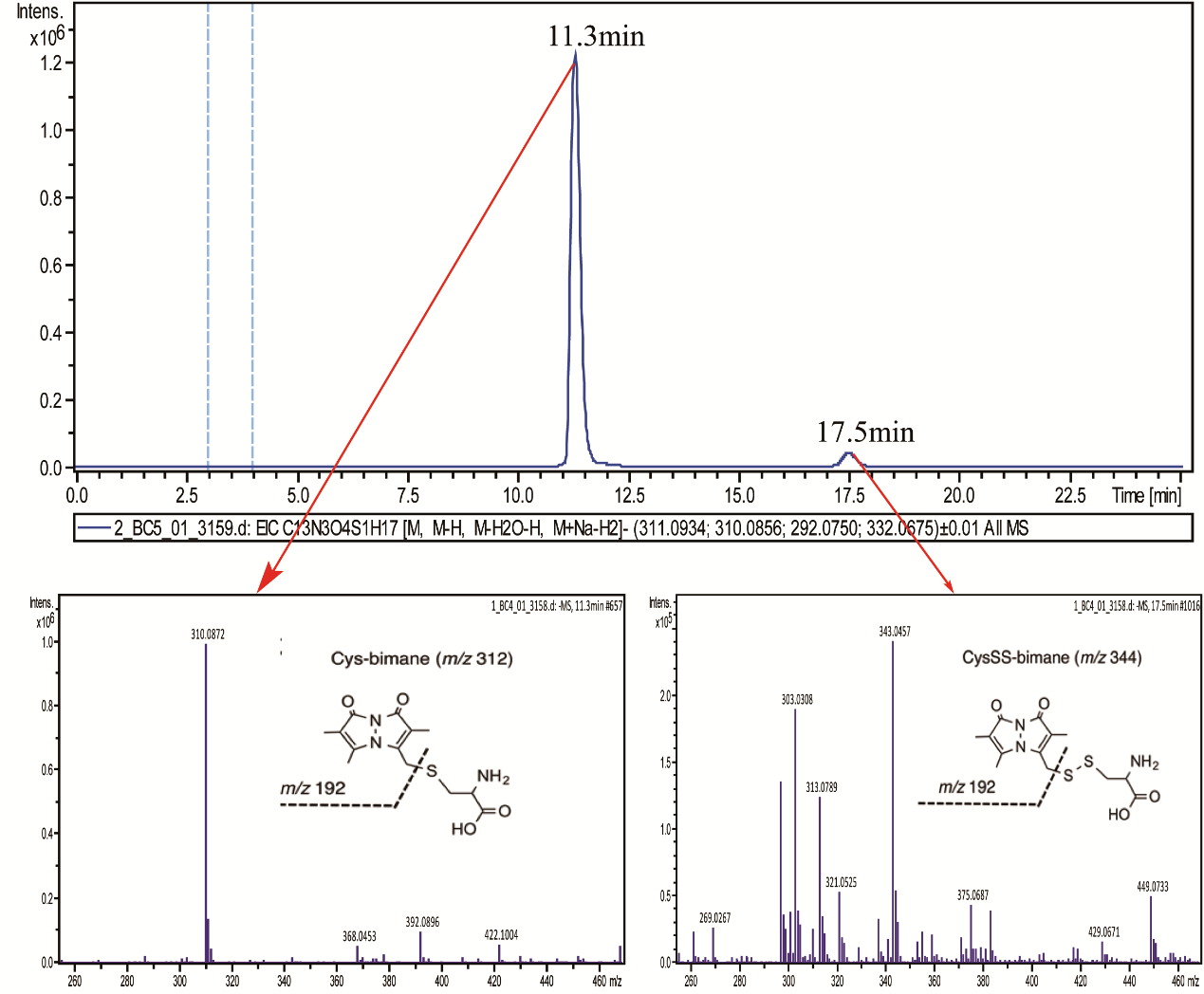

**Figure S1. LC-ESI-MS analysis of Crs1 produced RSS**. Crs1 (5 μM) was mixed with cysteine (100 μM) and PLP (100 μM) in reaction buffer (50 μM HEPES, 25 mM KCl, 15 mM MgCl2, pH 7.4). The reaction were conducted at 30^o^C for 30 min.

### Figure S2

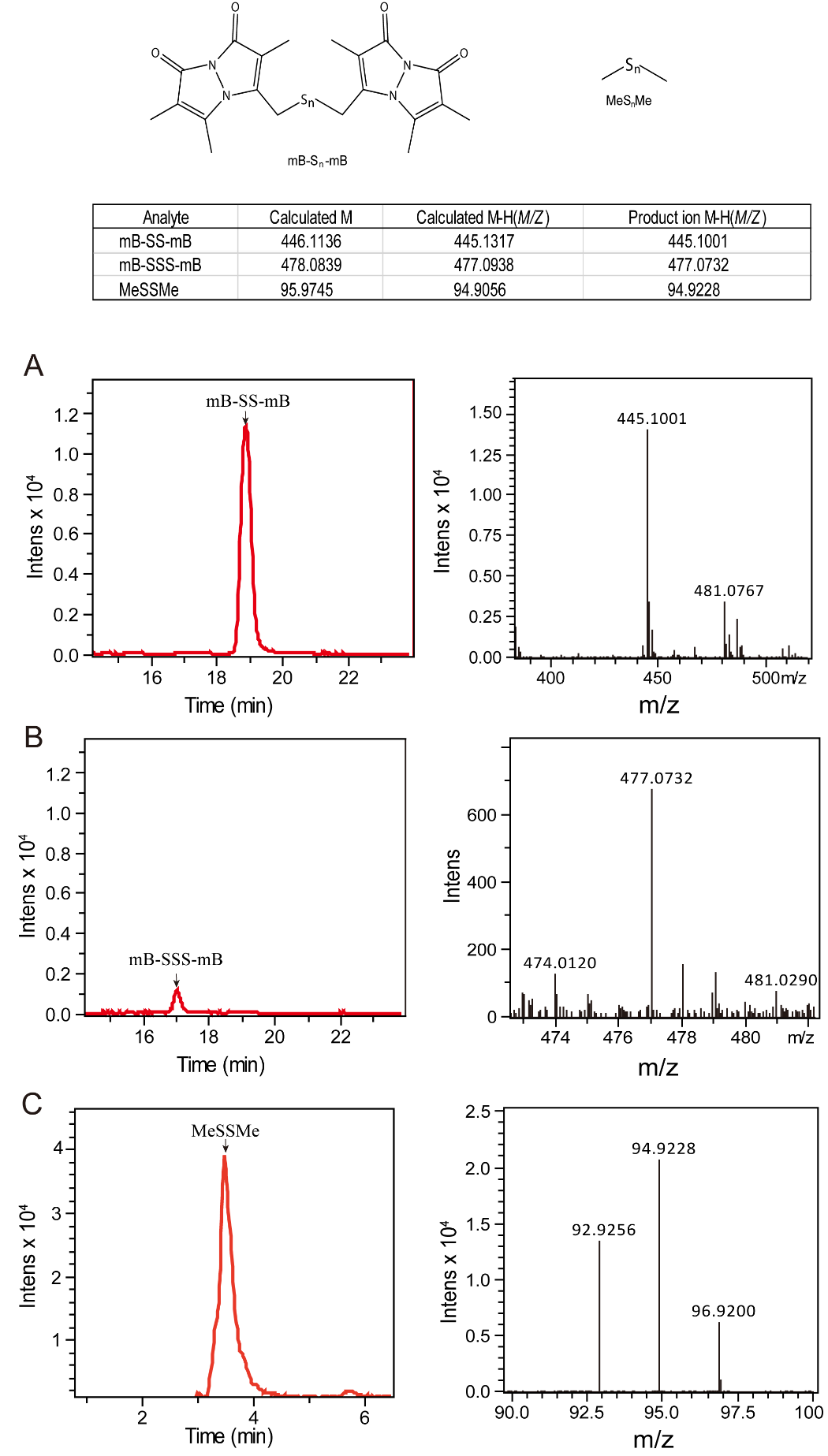

**Figure S2. LC-ESI-MS analysis of Rdl2 produced RSS.** Rdl2 (20 μg) were mixed with 5 mM thiosulfate (A and B) or MeSSSMe (C) in HEPES buffer (100 mM, pH 7.4). The reaction were conducted at room temperature for 30 min.

### Figure S3

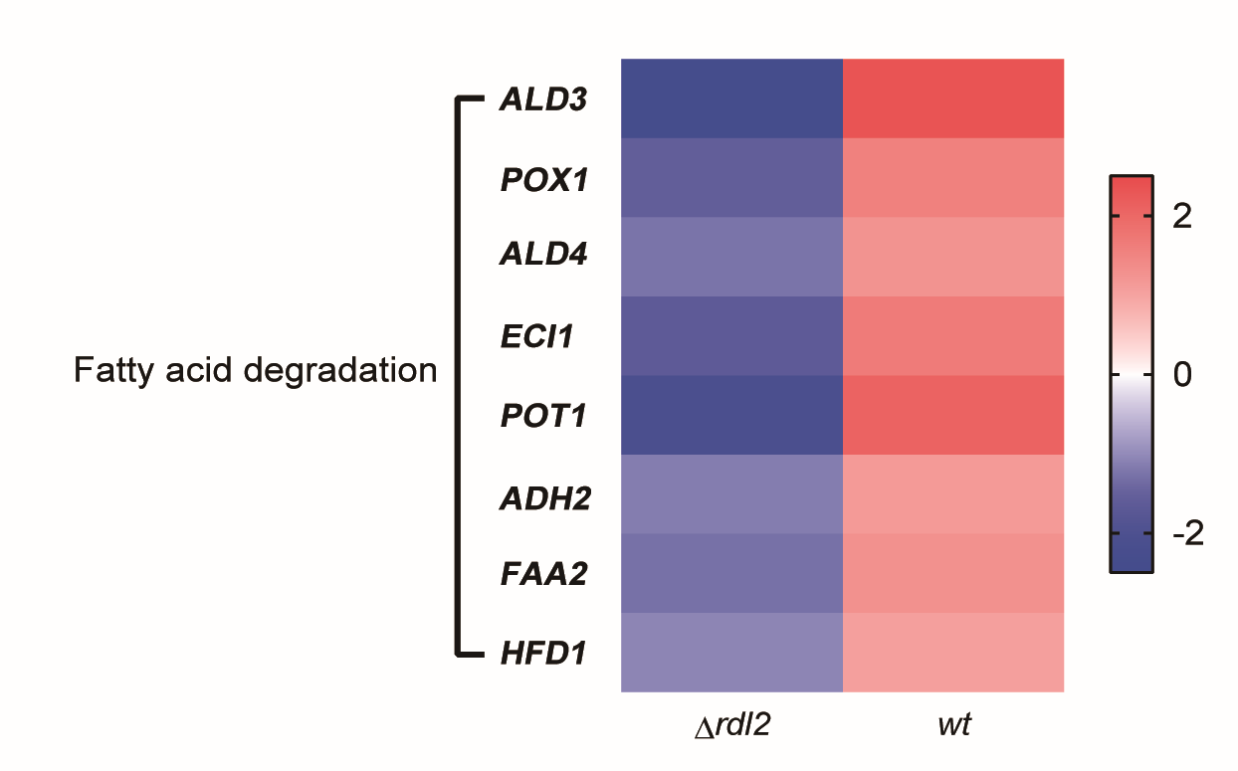

**Figure S3.** Transcriptional changes of genes related to fatty acid degradation.

### Figure S4

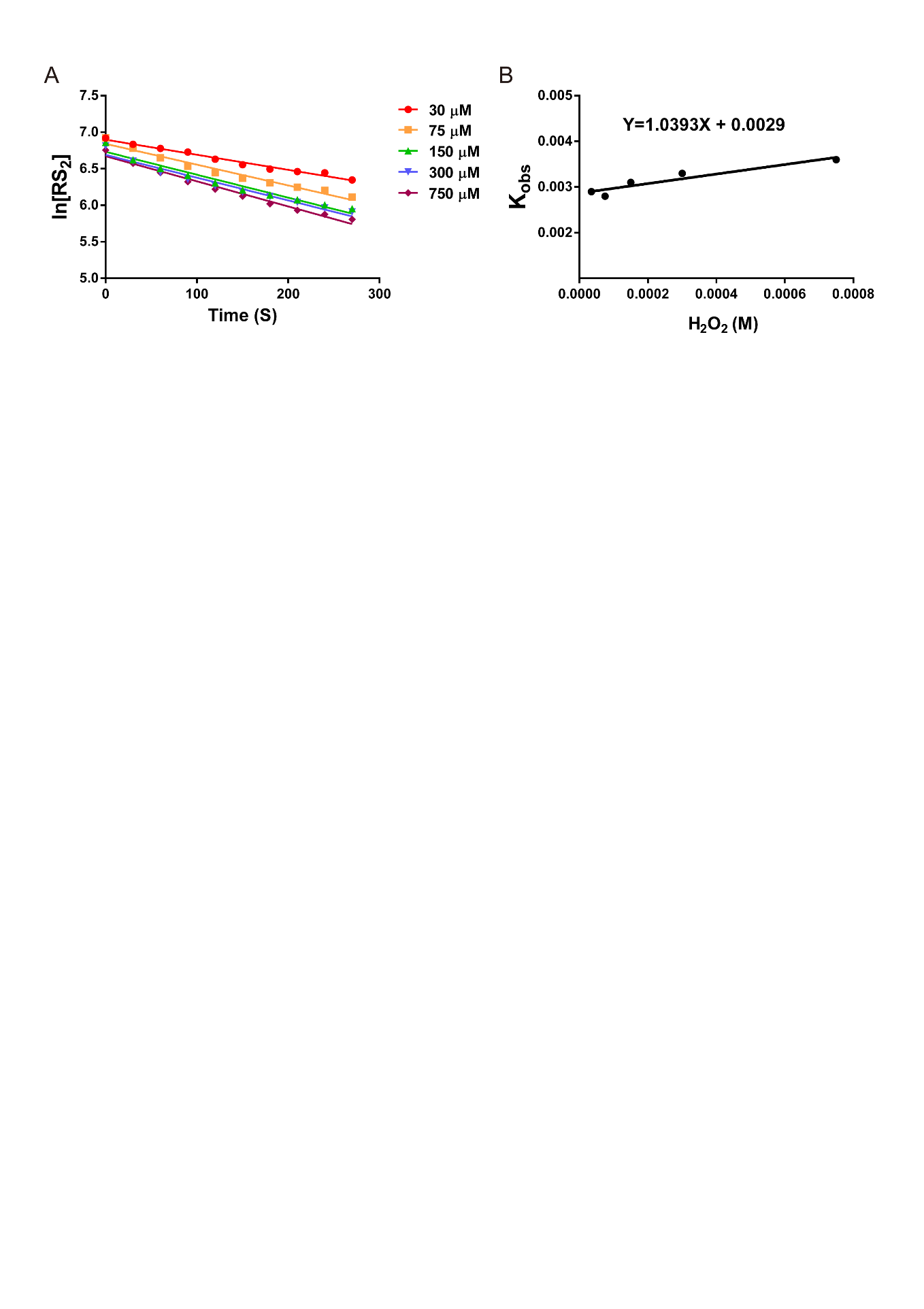

**Figure S4.** **Kinetic analysis of the reaction between S_8_ and H_2_O_2_.** A) The *k_obs_* value was calculated by plotting the *ln_[RS2]_* value against the reaction time. Colored lines represent *k_obs_* values obtained from different concentrations of H_2_O_2_. B) The 2^nd^ -order rate constant of the reaction was determined to be 1.04 M^−1^ s^−1^( *k_obs_* = *k* × *[H_2_O_2_]*). Data were from three independent repeats and shown as average ± s.d.

### Sheet 1. List of the targeted 200 metabolites

| **Metabolite name** | **Formula** |
| --- | --- |
| Deoxyinosine | C10H12N4O4 |
| Adenosine 3',5'-cyclic monophosphate (cAMP) | C10H12N5O6P |
| Glutathione disulfide | C20H32N6O12S2 |
| Nicotinamide adenine dinucleotide (NAD) | C21H27N7O14P2 |
| Betaine | C5H11NO2 |
| N6-Acetyl-L-lysine | C8H16N2O3 |
| Cytidine | C9H13N3O5 |
| D-Biotin | C10H16N2O3S |
| Adenosine | C10H13N5O4 |
| Cytidine 5'-monophosphate (CMP) | C9H14N3O8P |
| Adenosine monophosphate (AMP) | C10H14N5O7P |
| Inosine 5'-monophosphate (IMP) | C10H13N4O8P |
| Dihydroxy-acetone-phosphate | C3H7O6P |
| cis-Aconitic acid | C6H6O6 |
| Thiamine pyrophosphate (TPP) | C12H18N4O7P2S |
| Uridine diphosphate glucose (UDP-D-Glucose) | C15H22N2O18P2 |
| L-Homocysteine | C4H9NO2S |
| 5'-Methylthioadenosine | C11H15N5O3S |
| Deoxycytidine monophosphate (dCMP) | C9H14N3O7P |
| Uridine 5'-monophosphate (UMP) | C9H13N2O9P |
| Nicotinamide ribotide | C11H15N2O8P |
| S-Adenosylhomocysteine | C14H20N6O5S |
| Nicotinamide adenine dinucleotide phosphate (NADP) | C21H28N7O17P3 |
| Flavin adenine dinucleotide (FAD) | C27H33N9O15P2 |
| Acetyl-DL-Leucine | C8H15NO3 |
| S-Carboxymethyl-L-cysteine | C5H9NO4S |
| Dephosphocoenzyme A (Dephospho-CoA) | C21H35N7O13P2S |
| L-2-Hydroxygluterate | C5H8O5 |
| Deoxythymidine monophosphate (dTMP) | C10H15N2O8P |
| Deoxycytidine triphosphate (dCTP) | C9H16N3O13P3 |
| Guanosine diphosphate (GDP) | C10H15N5O11P2 |
| Thiamine monophosphate (TMP) | C12H17N4O4PS |
| Adenosine triphosphate (ATP) | C10H16N5O13P3 |
| D-glucosamine 1-phosphate | C6H14NO8P |
| Adenosine 5'-phosphosulfate (APS) | C10H14N5O10PS |
| Flavin mononucleotide (FMN) | C17H21N4O9P |
| Cytidine triphosphate (CTP) | C9H16N3O14P3 |
| Deoxyguanosine triphosphate (dGTP) | C10H16N5O13P3 |
| Uridine 5'-diphospho-glucuronic acid (UDP-D-Glucuronate) | C15H22N2O18P2 |
| N-Acetylputrescine | C6H14N2O |
| N-Carbamoyl-L-aspartic acid | C5H8N2O5 |
| Reduced nicotinamide adenine dinucleotide (NADH) | C21H29N7O14P2 |
| N-Acetyl-L-phenylalanine | C11H13NO3 |
| Tryptamine | C10H12N2 |
| L-2-Aminoadipic acid | C6H11NO4 |
| Phosphoenolpyruvate | C3H5O6P |
| Isobutyryl coenzyme A （Isobutyryl-CoA） | C25H42N7O17P3S |
| D-Glucuronic acid | C6H10O7 |
| Argininosuccinic acid | C10H18N4O6 |
| 2-keto-D-Gluconic acid | C6H10O7 |
| Glycyl-L-leucine | C8H16N2O3 |
| 4-Pyridoxic acid | C8H9NO4 |
| 2'-Deoxyguanosine 5'-diphosphate (dGDP) | C10H15N5O10P2 |
| Cholesterol sulfate | C27H46O4S |
| Phosphorylcholine | C5H14NO4P |
| sn-Glycerol 3-phosphate | C3H9O6P |
| Salicyluric acid | C9H9NO4 |
| Deoxyadenosine monophosphate (dAMP) | C10H14N5O6P |
| Biopterin | C9H11N5O3 |
| D-Neopterin | C9H11N5O4 |
| 4-Guanidinobutyric acid | C5H11N3O2 |
| Naringin | C27H32O14 |
| Pantothenic acid | C9H17NO5 |
| N-Acetylaspartylglutamic acid (NAAG) | C11H16N2O8 |
| 5'-Deoxyadenosine | C10H13N5O3 |
| Pyridoxal (Vitamin B6) | C8H9NO3 |
| 5-Methoxydimethyltryptamine | C13H18N2O |
| 3-Methyluric acid | C6H6N4O3 |
| Isobutyrylglycine | C6H11NO3 |
| N2,N2-Dimethylguanosine | C12H17N5O5 |
| 1-Methylxanthine | C6H6N4O2 |
| Alpha-N-Phenylacetyl-L-glutamine | C13H16N2O4 |
| Glycocholic acid | C26H43NO6 |
| D-Fructose 1,6-bisphosphate | C6H14O12P2 |
| N-Acetyl-L-alanine | C5H9NO3 |
| N6-methyladenosine | C11H15N5O4 |
| 2'-O-methyladenosine | C11H15N5O4 |
| N4-Acetylcytidine | C11H15N3O6 |
| Pseudouridine | C9H12N2O6 |
| 7-methylguanosine | C11H15N5O5 |
| 2-Thiocytidine | C9H13N3O4S |
| 3-Methyluridine | C10H14N2O6 |
| Acetylcholine | C7H15NO2 |
| Dimethylglycine | C4H9NO2 |
| D-Glucose 6-phosphate | C6H13O9P |
| Cytidine monophosphate N-acetylneuraminic acid | C20H31N4O16P |
| L-Anserine | C10H16N4O3 |
| Allantoic acid | C4H8N4O4 |
| 5-Aminolevulinic acid | C5H9NO3 |
| O-Succinyl-L-homoserine | C8H13NO6 |
| Sepiapterin | C9H11N5O3 |
| S-Lactoylglutathione | C13H21N3O8S |
| S-Nitroso-L-glutathione | C10H16N4O7S |
| N1-Methyl-2-pyridone-5-carboxamide | C7H8N2O2 |
| Hordenine | C10H15NO |
| GDP-L-fucose | C16H25N5O15P2 |
| Nicotinic acid adenine dinucleotide (NAAD) | C21H26N6O15P2 |
| gamma-L-Glutamyl-L-phenylalanine | C14H18N2O5 |
| gamma-Glutamyl-L-methionine | C10H18N2O5S |
| gamma-L-Glutamyl-L-valine | C10H18N2O5 |
| 5-Amino-4-carbamoylimidazole (AICA) | C4H6N4O |
| Fumaric acid/Maleic acid | C4H4O4 |
| Succinic acid | C4H6O4 |
| Nicotinic acid | C6H5NO2 |
| taurine | C2H7NO3S |
| Malic acid | C4H6O5 |
| Hypoxanthine | C5H4N4O |
| p-Hydroxybenzoic acid | C7H6O3 |
| 2,3-Dihydroxybenzoic acid | C7H6O4 |
| Orotic acid | C5H4N2O4 |
| Uric acid | C5H4N4O3 |
| myo-Inositol | C6H12O6 |
| Xanthurenic acid | C10H7NO4 |
| Uridine | C9H12N2O6 |
| Inosine | C10H12N4O5 |
| Sucrose | C24H40O5 |
| Ethanolamine | C2H7NO |
| Imidazole | C3H4N2 |
| Glycine | C2H5NO2 |
| gamma-Aminobutyric acid (GABA) | C4H9NO2 |
| Cytosine | C4H5N3O |
| Creatinine | C4H7N3O |
| Creatine | C4H9N3O2 |
| Nicotinamide | C6H6N2O |
| Thymine | C5H6N2O2 |
| L-Pipecolic acid | C6H11NO2 |
| L-Leucine | C6H13NO2 |
| L-Aspartic Acid | C4H7NO4 |
| Adenine | C5H5N5 |
| L-Glutamine | C5H10N2O3 |
| L-Glutamic acid | C5H9NO4 |
| L-Methionine | C5H11NO2S |
| L-Carnitine | C7H15NO3 |
| L-Phenylalanine | C9H11NO2 |
| L-Arginine | C6H14N4O2 |
| L-Citrulline | C6H13N3O3 |
| L-Tyrosine | C9H11NO3 |
| L-O-Phosphoserine | C3H8NO6P |
| N-Acetylglutamine | C7H12N2O4 |
| L-Tryptophan | C11H12N2O2 |
| N-Acetyl-D-glucosamine | C8H15NO6 |
| Deoxyguanosine | C10H13N5O4 |
| Guanosine | C10H13N5O5 |
| Riboflavin (Vitamin B2) | C17H20N4O6 |
| Sarcosine | C3H7NO2 |
| L-Lactic acid | C3H6O3 |
| Citraconic acid | C5H6O4 |
| Kynurenic acid | C10H7NO3 |
| Pyridoxine | C8H11NO3 |
| beta-D-Glucosamine | C6H13NO5 |
| L-Kynurenine | C10H12N2O3 |
| Flavone | C15H10O2 |
| Thiamine | C12H16N4OS |
| Folic acid | C19H19N7O6 |
| L-Proline | C5H9NO2 |
| L-Serine | C3H7NO3 |
| L-Valine | C5H11NO2 |
| Aminohippuric acid | C9H10N2O3 |
| N-Formylmethionine | C6H11NO3S |
| L-Dihydroorotic acid | C5H6N2O4 |
| L-Methionine sulfoxide | C5H11NO3S |
| L-Asparagine | C4H8N2O3 |
| L-Lysine | C6H14N2O2 |
| Glycerophosphocholine | C8H20NO6P |
| L-Homoserine | C4H9NO3 |
| Purine | C5H4N4 |
| Choline | C5H14NO |
| NG,NG-Dimethyl-L-arginine (ADMA) | C8H18N4O2 |
| L-Homocysteic acid | C4H9NO5S |
| S-Methyl-L-cysteine | C4H9NO2S |
| Xanthosine | C10H12N4O6 |
| Glyceric acid | C3H6O4 |
| beta-Hydroxybutyric acid | C4H8O3 |
| 5-hydroxy-Tryptophan | C11H12N2O3 |
| L-Abrine | C12H14N2O2 |
| Deoxycytidine | C9H13N3O4 |
| L-Carnosine | C9H14N4O3 |
| Dihydrothymine | C5H8N2O2 |
| N-Acetyl-L-tyrosine | C11H13NO4 |
| 2-Hydroxyadenine | C5H5N5O |
| Methylguanidine | C2H7N3 |
| DL-2-Aminooctanoic acid | C8H17NO2 |
| L-histidinol | C6H11N3O |
| Hydroxyisocaproic acid | C6H12O3 |
| Ornithine | C5H12N2O2 |
| histamine | C5H9N3 |
| Ureidopropionic acid | C4H8N2O3 |
| 7-Methylxanthine | C6H6N4O2 |
| Phenyllactic acid | C9H10O3 |
| Cytidine diphosphate choline (CDPcholine) | C14H26N4O11P2 |
| Urocanic acid | C6H6N2O2 |
| 5-Aminopentanoic acid | C5H11NO2 |
| N-Acetylcadaverine | C7H16N2O |
| Nicotinuric acid | C8H8N2O3 |
| 3,7-Dimethyluric acid | C7H8N4O3 |
| 6-Hydroxynicotinic acid | C6H5NO3 |
| cis-4-Hydroxy-D-proline | C5H9NO3 |
| 5-Methoxytryptamine | C11H14N2O |
| 2'-Deoxyguanosine 5'-monophosphate (dGMP) | C10H14N5O7P |
| Propionylglycine | C5H9NO3 |
